# A *number simplex* in the human medial temporal lobe

**DOI:** 10.64898/2026.06.25.734462

**Authors:** Hanlin Zhu, Assia Chericoni, Taha Ismail, Elizabeth A. Mickiewicz, Melissa C. Franch, Tarana Nigam, Xinyuan Yan, James Belanger, Ana G. Chavez, Jahnavi Nair, Danika Paulo, Eleonora Bartoli, Jay A. Hennig, Tomek Fraczek, Nicole R. Provenza, Seng Bum Michael Yoo, Hansem Sohn, Jessica F. Cantlon, Steven T. Piantadosi, Sameer A. Sheth, Benjamin Y. Hayden

**Affiliations:** Department of Neurosurgery, Baylor College of Medicine; Department of Electrical and Computer Engineering, Rice University; Department of Neuroscience, Baylor College of Medicine; Neuroengineering Initiative, Rice University; Department of Bioengineering, Rice University; Department of Biomedical Engineering, Sungkyunkwan University; Department of Psychology, Carnegie Mellon University; Department of Psychology, University of California Berkeley

## Abstract

Humans handle numbers nimbly, suggesting a richer neural manifold structure than the prevalent mental number line model. In populations of medial temporal lobe (MTL) neurons in humans performing two simple tasks (dot counting and arithmetic), we find robust neural coding of numerosity that results in high dimensional, **simplex-shaped** manifolds. This shape affords more flexibility than a linear manifold due to its high shattering dimensionality and expressibility. Dot arrays and Arabic numerals evoked distinct simplicial population codes, yet they were linked by a linearly transferable latent structure within the same task. We find similar simplicial geometry of number representations in large language models (LLMs). Moreover, subjects’ internally computed arithmetic results were decodable during the calculation period, with decoding accuracy correlating with individual mathematical capacity. Finally, linear transformations of simplicial operand representations modeled the brain’s conversion of operands into decodable results, suggesting that the brain’s arithmetic procedures have some resemblance to the attention architecture of LLMs. Together, these findings establish a high dimensional representational foundation for numerical cognition in the brain.

## INTRODUCTION

Humans have the ability to process numbers flexibly. We can understand a wide range of numbers, even ones we have never seen before, and represent them in multiple ways (Amalric & Dehaene, 2016; Dehaene et al., 2025). We use diverse strategies even for simple arithmetic, shifting among estimation, memory retrieval, decomposition, and rule-based procedures depending on the context (Banerjee et al., 2025; Campbell & Xue, 2001; Delazer et al., 2005; LeFevre et al., 2013; Siegler, 1988; Siegler & Braithwaite, 2017). We transition seamlessly between different numerical modalities (Piazza et al., 2007; Eger et al., 2003; Jacob & Nieder, 2009), and instantly grasp abstract, non-linear relationships between disparate quantities (Foucault et al., 2025). Even at a young age, we can skip-count by twos or threes, distinguish even from odd, or hold several nonconsecutive numbers (Gathercole et al., 2004) in working memory (e.g. PIN number, ZIP code).

Our mathematical flexibility implies the existence of a representational geometry capable of supporting the dynamic and combinatorial demands of symbolic human thought (Shepard et al., 1975; Dehaene et al., 2025). Indeed, classic multidimensional scaling behavioral studies suggest that the intrinsic representation of numbers may be high-dimensional (Shepard et al., 1975; Miller and Gelman, 1983; Griffiths and Kalish, 2002). These studies showed that people represent similarities among single-digit numerals not only by absolute magnitude difference but by multiple properties, including parity and arithmetic relatedness. At the neural level, this suggests a high-dimensional representational geometry.

The computational capacity of a neural population is strongly constrained by the geometry of its population-code. In particular, it depends on how many dichotomies of task conditions can be separated by a linear readout (shattering dimensionality), and on how flexibly those conditions can be mapped into output variables by downstream readouts (expressivity). High dimensional representations have high shattering dimensionality, making them ideal for on-the-fly flexible dichotomization (such as even/odd or prime/composite, Bernardi et al., 2020; Cover, 1965; Rigotti et al., 2013). They also have high expressibility, meaning they can support flexible reorderings (such as even-first, then-odd; Cover, 1967; Rigotti et al., 2013). In high dimensions, distinct population activity patterns can be arranged as the vertices of a high-dimensional generalization of a triangle or tetrahedron; such a shape is known as a *simplex* (Boyd & Vandenberghe, 2004). Because these vertices are affinely independent, simplicial geometry maximizes linear shattering of the finite pattern set and permits maximal linear-readout expressivity under the affine-readout assumptions (Cover, 1965). These facts motivated us to hypothesize that the inherent neural geometry of numerosity would involve a simplicial manifold structure.

Both human neuroimaging and non-human primate single-unit recordings have identified neural activity selective for specific numerosities (Nieder et al., 2002; Nieder et al., 2006; Harvey et al., 2013; Piazza et al., 2004). These classical “number neurons/voxels” typically exhibit Gaussian-like tuning curves centered on a preferred quantity, providing a direct biological substrate for the approximate number system and the behavioral distance effect. Neurophysiological investigations into numerical cognition have historically centered on the intraparietal sulcus and prefrontal cortex (Nieder & Dehaene, 2009; Debray et al., 2026). More recently, human single-neuron recordings have revealed robust number coding in the human medial temporal lobe (MTL), which includes the hippocampus and adjacent structures (Kutter et al., 2018, 2022, 2023, and 2024). This coding is consistent with emerging evidence that the MTL represents structured numerical information (Valério et al., 2026). While these results linking neural activity to number processing have generally been interpreted in the number line framework, they are also consistent with alternative structures because of the expressibility of high-dimensional manifolds.

The possibility of a high dimensional number manifold raises the question of how the brain implements arithmetic calculations. Pioneering studies using fMRI (Qin et al., 2014) and local field potentials (Pinheiro-Chagas et al., 2024) tell us *where* and *when* calculations occur, but not *how*. One important study identified abstract codes for arithmetic operation in the MTL (plus vs. minus, Kutter et al., 2022). Most recently, Czajko et al. (2024) investigated internally generated results during tasks involving the doubling or halving of dots on a screen with fMRI. However, this approach was limited to approximate, non-symbolic quantities and restricted the second operand to only two choices (two or four). Consequently, how the brain transforms two exact, symbolic operands into an exact result is unknown.

This problem is, ostensibly, more difficult with high-dimensional representations than with number lines. A similar challenge arises in large language models (LLMs), which operate over high-dimensional hidden representations (Zhao et al., 2026; Cheng et al., 2025) yet can successfully perform arithmetic computations (Stolfo et al., 2023). We investigated the hidden states of LLMs, finding that their numerical representations are organized as simplicial geometries. Inspired by this, we show that the human MTL also supports arithmetic processing through simplex-like symbolic representations, which flexibly integrate operand information into decodable internal results via operation-dependent linear transformations.

## RESULTS

Eleven patients undergoing intracranial monitoring for epilepsy (1 male, 10 females; age 41 ± 3.14 years old) participated in an arithmetic task. A subset (n = 10 patients) also performed a dot-counting task (**Figure 1A–C**; **Methods**). In the *<u>arithmetic task</u>*, participants viewed two operands (**Figures 1-3**) and an addition or subtraction operator, mentally computed the answer, and answered the question with a key press (**Methods**), followed by computer feedback (**Figure 1B**). In the *<u>dot-counting task</u>*, participants viewed an array of dots whose spatial layout varied across trials (Kutter et al., 2018), then reported the number of dots after a short delay (**Figure 1C**) with a key press, and then received feedback. Across both tasks, we recorded 554 isolated neurons from medial temporal lobe structures, including hippocampus (n = 389 neurons from 11 patients), entorhinal cortex (n = 70 neurons from 5 patients), amygdala (n = 77 neurons from 4 patients), and parahippocampal cortex (n = 18 neurons from 1 patient).

**Figure 1.**
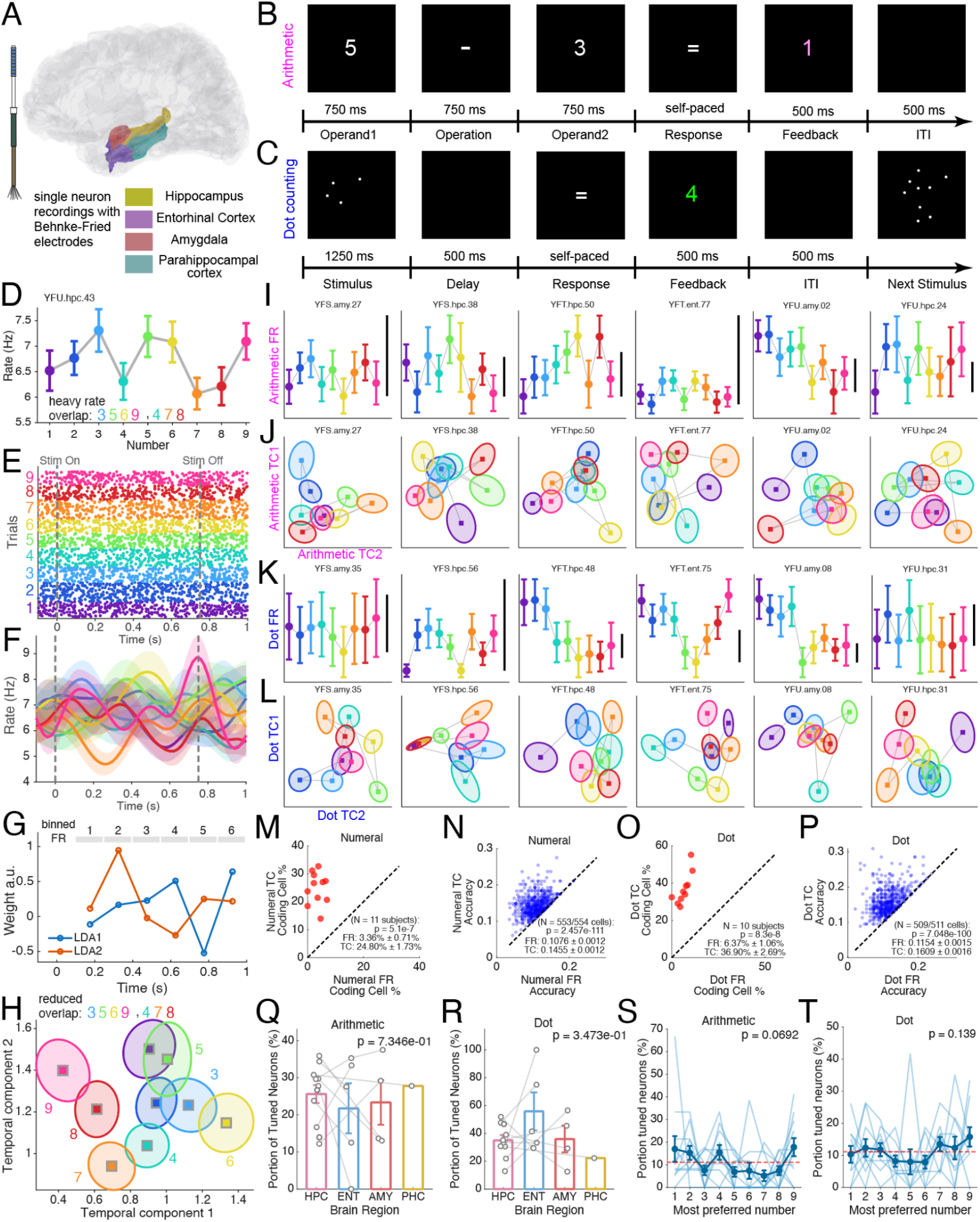
Temporally resolved single-neuron responses reveal number coding in the human MTL. **Abbreviations**: AMY, amygdala; ENT, entorhinal cortex; FR, firing rate; HPC, hippocampus; ITI, intertrial interval; PHC, parahippocampal cortex; TC, temporal component. **A**, Schematic of Behnke-Fried electrode recordings from medial temporal lobe regions included in the analysis. **B**, Arithmetic task structure. Participants viewed sequential arithmetic elements, entered a response, and received feedback; epoch durations are shown in the schematic. **C**, Dot-counting task structure. Participants viewed a dot array, reported the number of dots after a delay, and received feedback. **D**, Example neuron showing firing-rate tuning mean + SEM across number conditions during the arithmetic task. Colored numerals indicate conditions with overlapping mean firing rates. **E**, Spike raster from the same neuron in D, with trials grouped by number condition (color) and aligned to stimulus onset. **F**, Smoothed (Gaussian kernel *σ* = 150 ms) peristimulus firing-rate traces mean + SEM from the same neuron in E reveal number-dependent temporal dynamics despite overlap in mean firing rate. Colors indicate different numbers, as in D,E; numbers 7 and 9 showed activations/suppression of firing at specific timings that better distinguished them from others. **G**, Linear discriminant analysis applied to binned spike trains identifies temporal-component linear weights that discriminate number conditions across time. **H**, Projection of trials from the example neuron onto the first two temporal components reveals reduced overlap among number conditions. Squares indicate mean, and ellipses indicate covariance normalized by square root of trial count, matching the mean + SEM (standard deviation normalized by square root of trial count) in D. **I**, Same as D for additional example neurons during the arithmetic-task. Different numbers give overlapping firing rates. Black scale bar = 1 Hz. **J**, Same as H, temporal-component projections for the same neurons in I reveal clearer separation of number conditions at the single-neuron level. **K**, Same as I but for neurons in the dot-counting task. **L**, Same as J but for temporal-component projections of the neurons in K. **M**, Subject-level arithmetic coding-cell proportions under firing-rate and temporal-component definitions. The dashed line indicates equal coding-cell proportion. **N**, Cross-validated arithmetic-task decoding accuracy for firing-rate and temporal-component based representations. Each point represents one neuron, and the dashed line indicates equal performance, N = 553 neurons with firing rate greater than 0.007 Hz during the task out of the total 554 neurons. **O**, Subject-level dot-counting coding-cell proportions under firing-rate and temporal-component definitions. The dashed line indicates equal coding-cell proportion. **P**, Cross-validated decoding accuracy of quantity during dot-counting task for firing-rate and temporal-component representations. Each point represents one neuron, and the dashed line indicates equal performance. N = 509 neurons with firing rate greater than 0.007 Hz during the task out of the total 511 neurons. **Q**, Proportion of tuned neurons (temporal-component definition) across medial temporal lobe regions in the arithmetic task. Dots indicate individual subjects and linked lines indicate the same subject (N = 11), plotting mean + SEM. **R**, Same as in Q but for the dot-counting task. N = 10 subjects. **S**, Distribution of most preferred arithmetic-task numbers among numeral coding neurons. Light lines indicate individual subjects, dark markers indicate mean + SEM, and the dashed line indicates chance. **T**, As in S but for the dot counting task.

Neural tuning for task variables is often quantified using a standard fixed-window firing-rate code. However, a good deal of previous research indicates that much of the informational capacity of neuronal firing could reside in the spiking temporal dynamics beyond their averaged firing rates (Cariani & Baker, 2025; Li et al., 2026; Panzeri et al., 2026; She et al., 2024; Smith et al., 2019; Xie et al., 2024; Zuo et al., 2015; Chabrol et al., 2015). Indeed, in an example neuron during the arithmetic task, firing-rate tuning curves showed overlapping responses across different numbers (**Figure 1D**). Yet, in the same example neuron, spike rasters and peristimulus time histograms revealed that numbers with overlapping mean firing rates nevertheless produced different response dynamics over time (**Figure 1E,F**), consistent with temporal pattern separation (Madar et al., 2019a, 2019b; Nigam et al., 2025). We therefore binned each trial’s spike train across time and used cross-validated linear discriminant analysis (LDA) to extract stimulus-coding temporal components, defined as linear combinations of time bins (Zhu et al., 2025, **Figure 1G**). Projecting trials from the example neuron onto the first two temporal components revealed additional details of the number code (**Figure 1H**). Note that we specifically confirmed that our temporal component extraction procedure does not artifactually generate high dimensional representational geometry (**Supplementary Figure 1**).

This pattern generalized across neurons and tasks. In six example neurons during the arithmetic task, firing-rate tuning curves often showed overlapping responses across different numbers (**Figure 1I**), whereas the corresponding temporal-component projections separated number conditions more clearly at the single-neuron level (**Figure 1J**). We observed a similar pattern in the dot-counting task: example neurons showed partially overlapping firing-rate tuning, but temporally resolved projections separated dot-number responses along one or more temporal components (**Figure 1K, L**).

To quantify the increase in sensitivity of incorporating temporal coding, we compared cross-validated nine-way number decoding using both firing rate and temporal-component representation. The proportion of number-coding neurons in the arithmetic task increased from 3.36 ± 0.71% to 24.80 ± 1.73% under the temporal code (paired t-test across n = 11 subjects, p < 0.0001; significant neurons defined from permutation test with label shuffling, **Figure 1M**). Across all neurons, temporal components significantly improved single-neuron decoding accuracy relative to firing rate by 3.79% (**Figure 1N**). Temporal dynamics also produced a more sensitive measurement of neural coding in the dot-counting task. The proportion of numerosity (non-symbolic number)-coding cells increased from 6.37 ± 1.06% under the firing-rate code to 36.90 ± 2.69% under the temporal code across subjects (paired t-test, n = 10 subjects, p < 0.0001; **Figure 1O**). Similarly, across n = 509 neurons, incorporating temporal components increased single-neuron decoding accuracy from 0.1154 ± 0.0015 to 0.1609 ± 0.0016 (paired t-test, p = 7.048e-100; **Figure 1P)**. Thus, for both symbolic arithmetic stimuli and nonsymbolic dot stimuli, temporally resolved single-neuron responses revealed substantially more information than firing rate alone.

We found no significant regional differences in the proportion of neurons coding numbers in each MTL region. Specifically, using the temporal-component tuning definition, we fit a binomial generalized linear mixed-effects model to explain the proportion of coding neurons with brain region as a fixed effect and subject as a random effect, allowing us to test regional differences while accounting for repeated neurons sampled from the same patient. The post-hoc ANOVA region effect was non-significant in both the arithmetic task (region main effect: F(3,550) = 0.43, p = 0.74; **Figure 1Q**) and the dot-counting task (region main effect: F(3,507) = 1.1034, p = 0.35; **Figure 1R**). These results are consistent with others showing that number-coding neurons are widely distributed across the recorded medial temporal lobe regions (Kutter et al., 2018; 2024).

Finally, we tested whether the observed coding neurons were dominated by any particular especially well-represented numbers (cf. Kutter et al., 2024). For each coding neuron, we defined the most preferred number as the number with the highest decoding accuracy, then compared the distribution of preferred numbers across subjects. In the arithmetic task, we observed no evidence for favored numbers across 1–9 after accounting for subject-level variability (Friedman test, p = 0.0692; **Figure 1S**). Thus, while there is a marginal trend suggesting a preference for numbers at the high and low end of the range tested (numbers 1 and 9) across neurons (Kutter et al., 2024), it is not significant in our current data. The dot-counting task showed the same pattern, with no significant overrepresentation of any individual preferred number (Friedman test, p = 0.139; **Figure 1T**).

### A simplicial number manifold in MTL

We next investigated the shape of the neural population manifold of numbers in MTL. A number line, even when log-spaced, imposes a privileged ordering and supports a limited set of dichotomies and orderings. In contrast, a simplex-like (“*simplicial*”) geometry supports many dichotomies and orderings (**Figure 2A,B**). In an example subject, z-scored temporal components pooled across coding neurons showed structured number tuning, with components preferring a given dot number often also responding to other dot numbers, not necessarily only to the immediate neighbouring numbers (**Figure 2C**). However, the centroids of numbers 1-9 did not fall along a line; non-adjacent numbers did occupy nearby positions, suggesting a simplicial population geometry (**Figure 2D**).

**Figure 2.**
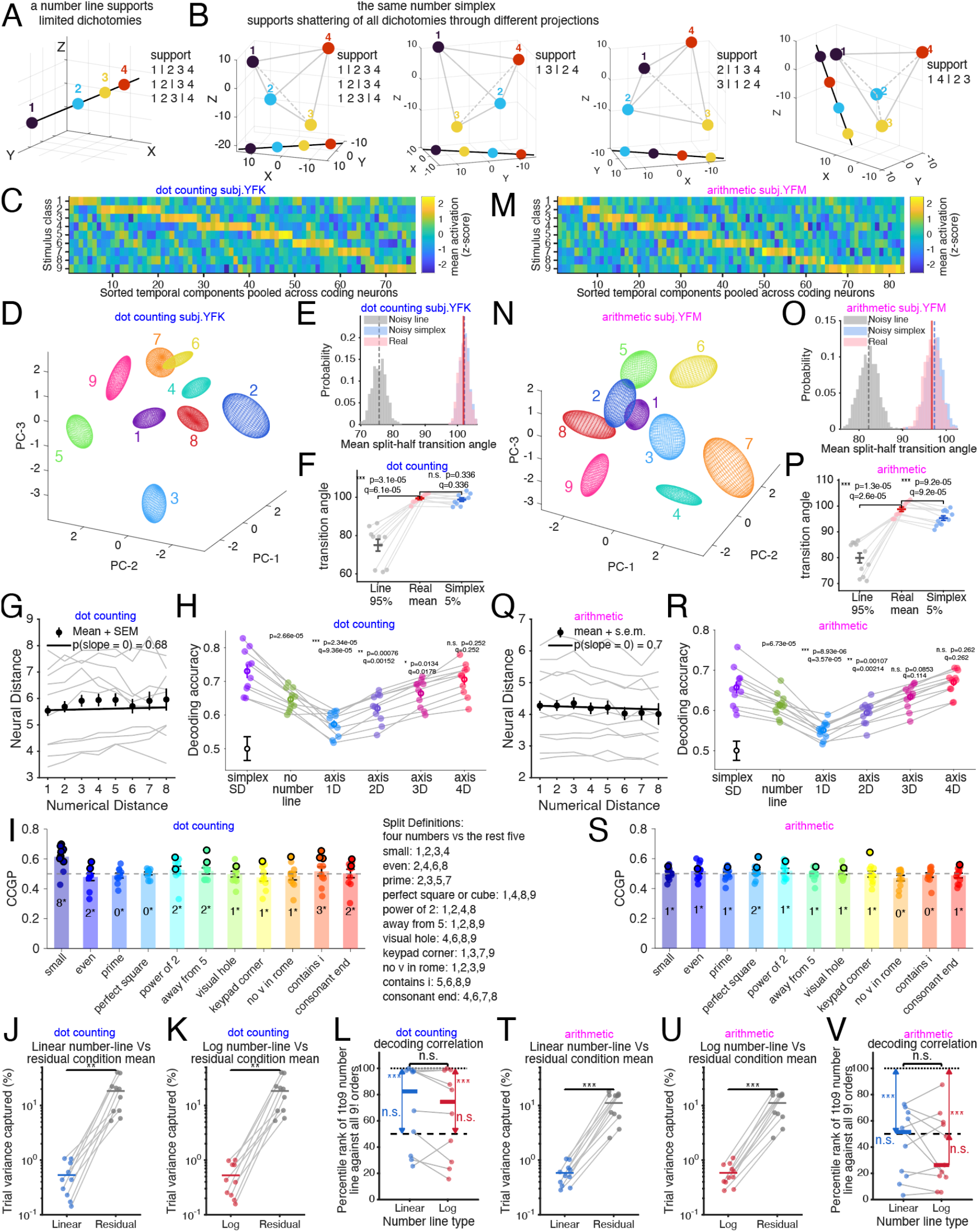
Medial temporal lobe number populations form high-dimensional simplex-like manifolds rather than a privileged one-dimensional number line. **Abbreviations**: CCGP, cross-condition generalization performance; MTL, medial temporal lobe; PC, principal component; SD, shattering dimensionality; SEM, standard error of the mean. **A**, Schematic illustrating that a one-dimensional number line supports only limited dichotomies through partition hyperplanes. **B**, Schematic illustrating that the same set of number identities arranged as a simplex can support many dichotomies through different linear projections onto a line. **C**, Example dot-counting subject showing z-scored mean activation of sorted temporal components pooled across coding neurons. Rows: dot-number stimulus classes; columns: temporal components sorted by preferred class and then the second most preferred class. **D**, Three-dimensional principal-component projection of dot-counting population activity (z-scored pooled temporal components across tuned neurons) for an example subject. Colored ellipsoids (mean + 0.5*covariance) indicate number-specific activity distributions, showing that dot-number centroids do not lie on a single ordered line. **E**, Split-half transition-angle distribution for the dot-counting example subject compared with matched noisy number-line and noisy-simplex controls. Vertical lines indicate means. This analysis was done in the unreduced, high dimensional neural space. **F**, Subject-level dot-counting transition-angle comparison among the noisy-line control, empirical data, and matched-simplex control. Gray lines connect different conditions of the same subject; transition angles were calculated from arccos function and were restricted to 0 to 180 degrees when multiple angles satisfy the same cosine (normalized vector dot product) value. **G**, population neural distance (Euclidean) as a function of numerical distance. Gray: individual subjects; black: mean ± SEM across subjects. Black line: fitted linear mixed effect slope. **H**, shattering dimensionality during the dot-counting task. Performance is shown for the full representation, magnitude-residualized representation (no number line), and PCA-restricted subspaces. Gray lines connect subjects, colored points indicate subject-level values, and black markers indicate a label-shuffled null distribution, 160 shuffles per subject, pooled across subjects, the hollow black circle indicating mean and whiskers indicate 2.5% to 97.5%. Statistics above each column indicates paired t-test results against the full population SSD (first column). **I**, Dot-counting cross-condition generalization performance for interpretable numerical, visual, keypad, and verbal dichotomies. Bars show CCGP across subjects in mean + SEM, dots show individual subjects, the dashed line marks chance, and numbers above bars indicate the number of subjects with significant generalization at p < 0.05; Central inset, Definitions of the interpretable dichotomies used for the cross-condition generalization analyses in panels I and S. **J**, Dot-counting variance captured by the best-fitting linear number-line projection compared with residual condition-mean structure after removing that projection. **K**, Same as panel J, but for a logarithmic number-line projection. **L**, Dot-counting single-trial regression based decoding analysis, comparing the rank of the canonical 1-to-9 ordering with all possible ordered number lines for both linear (blue) and logarithmic (red) number-line definitions. Dashed lines indicate chance rank (thick) and the best possible rank (thin). **M-V**, Same as panels **D-L**, but for Arabic numerals in the arithmetic task.

To quantify this population geometry, we first measured transition angles between each pair of sequential numbers (Hénaff et al., 2019; 2021). A number line predicts transition angles near 0°, whereas a noiseless regular simplex predicts transition angles of 120° (e.g., the 120° turn required to navigate from vertices 1 to 2, and then 2 to 3, along the edges of an equilateral triangle, which is a regular 2-simplex; this also holds for regular simplices in higher dimensions due to the equidistance property). Because trial noise can inflate transition angles even for a line-like code, we constructed split-half distributions and compared them with two matched controls: (1) a log-number-line control in which number centers were shifted onto a fitted log line while preserving trial-to-trial noise, and (2) a matched simplex control in which centers were placed on an 8-dimensional simplex with distance from the center to vertices scaled to match the empirical distances. In the same example subject, the empirical transition-angle distribution was far from the number-line control and largely overlapped the matched simplex control (**Figure 2E**). Across subjects, the empirical mean transition angle was significantly greater than the 95th percentile of the number-line null distribution, indicating that the geometry was substantially more simplicial than expected from a noisy number line (**Figure 2F**). In contrast, the empirical transition angles were not significantly different from the lower bound of the matched simplex distribution, consistent with a simplex-like geometry rather than a noise-exaggerated one-dimensional line (**Figure 2F**).

We next asked whether neural distances recapitulate numerical distances, as expected under a number-line representation. We fit a mixed-effects model predicting neural distance from numerical distance, with subject and number identities included as random effects to account for dependence among entries in the representational distance matrix. Numerical distance did not significantly predict neural distance in the dot-counting task across patients (slope β = 0.014735; p[slope = 0] = 0.68; **Figure 2G**). Thus, although the population clearly distinguished number identities, neural distances did not reliably scale with numerical distance, providing no evidence for a smooth number-line-like metric.

### High Shattering Dimensionality

One benefit of a simplicial code is high shattering dimensionality, which refers to the number of classification dichotomies available (Bernardi et al., 2020; Posani et al., 2025). We quantified simplex shattering dimensionality (SSD) by decoding dichotomies across all possible number splits (rather than restricting the analysis to balanced half-splits). The full population produced high shattering performance (decoding accuracy or SSD = 0.7302 ± 0.0203; n = 10 subjects.

**Figure 2H**). Critically, all subjects’ SSD values were above chance, thus no single patient’s representation is purely one-dimensional (p = 0.0062, permutation test with 160 shuffles). Importantly, this dimensionality was not explained simply by residual magnitude or number-line information: shattering performance remained significant even after regressing out magnitude-related structure (p = 0.0062, **Figure 2H**), despite being expectedly smaller than the full population (p = 0.000026 two-sided paired t-test). To estimate the minimum dimensionality required to support this performance, we projected the number manifolds into lower-dimensional PCA subspaces and recomputed the shattering score. One-, two-, and three-dimensional projections all produced significantly lower shattering performance than the full representation or high-dimensional reference (1D: p = 2.34e-5, q = 9.36e-5; 2D: p = 7.6e-4, q = 0.00152; 3D: p = 0.0134, q = 0.0178, q denotes the Benjamini-Hochberg-corrected p-value), whereas the four-dimensional projection was no longer significantly different from the original, unreduced high dimension neural code (4D: p = 0.252, q = 0.252; **Figure 2H**). Thus, while the true dimensionality is likely higher, four dimensions serve as a conservative lower bound necessary to account for the observed shattering dimensionality given our data.

### Low CCGP

We next asked whether the number manifold has specific abstract axes that are strong enough that they are recoverable across novel contexts (like a “baked-in” representation of even-odd), or whether these categories are flexibly created on the fly. We therefore selected 11 meaningful dichotomies of the nine numbers: (1) magnitude, (2) visual, (3) keypad, and verbal groupings, and measured cross-condition generalization performance (CCGP) across subjects (**Figure 2I**). For each dichotomy, we trained a linear classifier to separate the categories using a subset of the numbers, and tested its accuracy on the remaining, held-out numbers (e.g., training a “small vs. large” axis on {1, 3, 4} vs. {5, 6, 7, 9} and testing generalization on {2} vs. {8}). Consistent with a high-dimensional representation, CCGP was generally weak (Bernardi et al., 2020): except for the “small/large” split, which generalized in 8 of 10 subjects, most other dichotomies showed significant generalization in only a few (0–3) subjects. In other words, aside from the high-low magnitude split, these other dichotomies do not form highly abstract, reusable axes within the manifold that could be linearly read out across novel contexts.

Mean CCGP across splits was 0.5096 ± 0.0076, with per-split values ranging from 0.3037 to 0.6976, where a CCGP of 0.5 is chance level. Collectively, the pairing of high shattering dimensionality with low CCGP reflects a high-dimensional neural code optimized for memorization over rule-based abstraction (Bernardi et al., 2020). While the high shattering dimensionality reflects the capacity to separate highly familiar digits into many arbitrary groupings, the weak CCGP indicates that the manifold lacks the abstract axes required to rapidly categorize novel unseen numbers (e.g. a newly discovered (fictional) integer quantity between four and five) into these learned groupings.

### No privileged representation of ordered number line

Whereas the CCGP asks whether binary categorical axes generalize across held-out numbers, we next used a complementary variance-decomposition analysis to ask how much of the reliable number-condition structure is carried by a continuous ordered number-line axis. To establish the amount of reliable condition-dependent structure available to be explained, we replaced each trial’s population response with the mean population response for its corresponding number condition. This condition-mean representation removes trial-by-trial variability while preserving stable differences among number conditions. Across subjects, this reliable condition-mean structure accounted for approximately 19% of the total trial-level variance. We then fitted a 1-9 number-line projection using the individual single-trial responses through linear ridge regression and evaluated how much of the condition-mean structure was captured by projecting the condition means onto this axis. If the number conditions were primarily arranged along an ordered one-dimensional continuum, this projection should capture most of the condition-mean variance, leaving little residual condition-mean structure. Instead, the number-line component accounted for only a small fraction of the reliable signal. The best-fitting linear number-line projection explained 0.5330 ± 0.1124% of total trial variance, whereas the residual condition-mean structure left outside the number-line component explained 18.4155 ± 4.1041% (p = 0.0018, two-sided paired t-test; **Figure 2J)**. A similar result was obtained removing a logarithmic version of the number-line (log number line: 0.5267 ± 0.1078% variance explained; residual condition mean: 18.4218 ± 4.0881%; p = 0.0017; **Figure 2K**). Separately, at the single-trial level, we trained cross-validated regression models to project neural activity onto each of the 9! possible ordered lines and ranked the canonical 1–9 ordering by held-out decoding correlation. The canonical ordering did not rank significantly above chance (1-9 line vs 50%, linear: p = 0.0527; log: p = 0.0967, one-sided Wilcoxon signed rank test) and remained significantly below the best possible ordering (1-9 line vs 100%: p = 9.77e-04; log vs 100%: p = 9.77e-04; **Figure 2L**). No difference was found between the ranking of linear vs log number line (p = 0.19). Thus, although a number-line projection can always be extracted from a high-dimensional code, it is not a privileged axis of the neural representation.

### A simplicial manifold for Arabic numerals

We repeated the same analyses for Arabic numerals in the arithmetic task (example: **Figure 2M-P)**. As in dot counting, we found that numerical distance did not predict neural distance across the representational distance matrix (**Figure 2Q**), and concomitant high shattering dimensionality (**Figure 2R**) and a minimal dimensionality of three (**Figure 2R**). We speculate that the lowered bound on estimated dimensionality for Arabic numerals may reflect the reduced stimulus variability (geometry of the dot locations) of symbolic numerals compared with spatially variable dot arrays. CCGP across the same 11 dichotomies was limited (**Figure 2S**). The linear and log number-line projections remained weak relative to the full condition-mean structure (**Figure 2T-U**). The canonical 1–9 order was not uniquely privileged at the cross-validated single-trial decoding level. The canonical ordering (**Figure 2V**) did not rank significantly above chance (linear: p = 0.77; log: p = 0.91, one-sided Wilcoxon signed rank test) and remained significantly below the best possible ordering (linear vs 100%: p =4.88e-04; log vs 100%: p =4.88e-04). No difference was found between the ranking of linear and log number line (two-sided Wilcoxon signed rank test, p = 0.52).

### Arabic and dot numbers have different but related codes

We next asked whether MTL number representations are shared across nonsymbolic and symbolic formats. We took advantage of the Arabic-numeral feedback period in the dot-counting task. This allowed us to compare neural codes for nonsymbolic dot numerosities and symbolic Arabic numerals within the same counting-task context.

At the single-neuron level, dot-stimulus and numeral-feedback showed limited preference consistency. The number of neurons with correlated cross-format number decoding accuracy was not more frequent than expected by chance (p = 0.77, two-sided binomial test at a = 5%, **Figure 3A**): only 5.2% of neurons that significantly coded number in both epochs showed similar numerical preferences. Consistent with this, the correlation between most preferred dot number and most preferred feedback-numeral number was not significant across neurons (r = 0.09, p = 0.501; **Figure 3B**). Thus, individual MTL neurons generally did not express a strictly format-invariant number preference, even when dots and Arabic numerals appeared within the same task.

**Figure 3.**
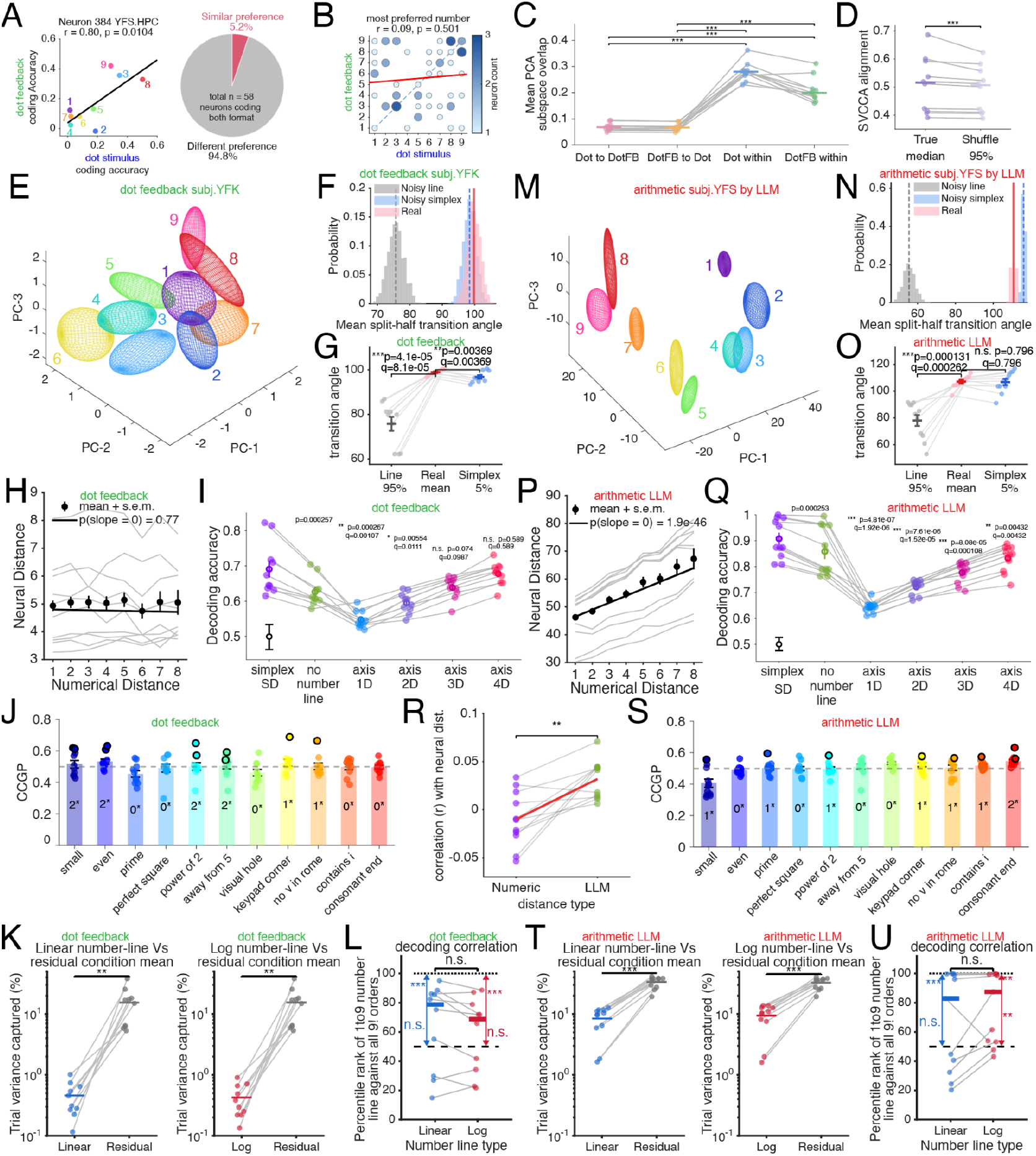
Different yet linked number representations between dot and numeral formats. **A**, Left: Example neuron coding both dot-stimulus numerosity and Arabic-numeral feedback coding within the dot-counting task. Each point shows the coding accuracy for one number across the two formats; colors indicate number identity, and the black line shows the best linear fit; Right: Fraction of neurons coding both dot-stimulus and numeral feedback formats that showed similar number preference across formats. **B**, Preferred-number matrix for neurons coding both dot-stimulus and numeral feedback formats. Circle size and shading indicate neuron count, the dashed diagonal indicates identical preferred numbers across formats, and the red line shows the linear trend. **C**, Population subspace overlap between dot-stimulus and numeral feedback representations. Cross-format overlap is compared with split-half within-format overlap for each format; gray lines indicate subjects, colored markers show per subject values or means, and brackets indicate paired statistical comparisons. **D**, SVCCA alignment between dot-stimulus and dot feedback numeral population codes compared with a label-shuffled control. Gray lines indicate subjects, and points show true median alignment and the shuffled-control 95th percentile. **E-L**, Same as Figure 2 D-L but for numeral representation during the feedback period of the dot counting task. **M-U**, Same as Figure 2 N-V but for numeral representation during the arithmetic task in LLM hidden-state embeddings. **R**, Subject-level comparison of correlations between MTL arithmetic neural distance structure and either numerical-distance structure or LLM representational-distance structure. Gray lines connect paired subjects, and the ends of red line segments indicate the group means.

However, unified cognitive variables need not be represented by identical tuning at the level of single neurons. They can instead be embedded within the shared low-dimensional geometry of a broader population response (Gallego et al., 2018; Fine et al., 2023; Safaie et al., 2023). We therefore asked whether dot stimuli and feedback numerals occupied similar number-related population subspaces. For each subject and format, we computed the principal-component subspace spanned by number-related population activity. We then quantified subspace overlap by asking how much variance from one format was captured by the principal components of the other format (Elsayed et al., 2016; Hu et al., 2026; Yan et al., 2026a). Within-format overlap was estimated using repeated split-half resampling, providing a reliability-matched upper bound for the overlap expected given finite trial counts.

This analysis initially suggested strong format dependence at the population level. Cross-format overlap between dot stimuli and dot-feedback numerals was substantially lower than within-format overlap (p < 0.001, one-sided Wilcoxon signed-rank test; **Figure 3C**). Thus, dot and feedback-numeral representations did not simply occupy the same neural dimensions.

Low subspace overlap, however, does not rule out the possibility that the two formats use different axes that are nevertheless linearly transformable through a shared latent code (Gallego et al., 2020). We therefore applied singular vector canonical correlation analysis (SVCCA) to test whether dot and feedback-numeral representations could be aligned after allowing linear rotations and rescaling of their activity patterns (Raghu et al., 2017). In this analysis, a high SVCCA score would indicate that two formats express similar latent numerical structure, even if that structure is read out through different population axes.

Unlike the low subspace overlap, SVCCA revealed significant alignment between dot-stimulus and feedback-numeral representations. The true dot–feedback alignment exceeded the 95th percentile of a label-shuffled control distribution (true median alignment: 0.5155 ± 0.0313; shuffle 95%: 0.5070 ± 0.0261; median ± s.e.m.; p = 9.77e-4, one-sided Wilcoxon signed-rank test; **Figure 3D**). Thus, within a matched task context, dot stimuli and Arabic feedback numerals could be expressed in different population subspaces while still preserving a linearly recoverable shared numerical structure. In other words, format-dependent neural codes can nevertheless reflect a common latent number geometry when the broader task demands are held constant.

Interestingly, applying the same analyses to dot stimuli and Arabic numerals during the arithmetic trials did not reveal such linear alignment (**Supplementary Figure 3**). Thus, the shared latent structure observed for dot stimuli and feedback numerals depends critically on matching the task context, not merely on comparing representations of the same numerical values (see **Discussion**).

### Simplicial manifold for Arabic numbers in feedback period of the dot task

Arabic numbers shown during the dot-counting task feedback period exhibited simplicial geometry. Specifically, the transition angle distribution was far larger than expected under a noisy number line control and fell within the range of the simplex (**Figure 3E-G**). The distance between neural evoked responses did not correlate with numerical distance (**Figure 3H**), the resulting manifold had high shattering dimensionality (**Figure 3I**), cross condition generalization for feedback numbers was limited (**Figure 3J**), and both linear and log number-line projections explained only a small fraction of trial variance compared with the residual condition-mean structure left after removing the number-line component (**Figure 3K**). Indeed, the canonical 1–9 ordering was not consistently ranked above chance among all possible ordered lines and was significantly below the best possible ordering (**Figure 3L**).

### Simplicial geometry in LLM embedding captures MTL neural representations better than numerical distance

LLMs represent a kind of model organism that allows for direct comparison with representations of structured information in a non-brain system (Goldstein et al., 2025; Doerig et al., 2025; Caucheteux et al., 2023; Schrimpf et al., 2021; Franch et al., 2026a, 2026b; Yan et al., 2026a, 2026b; Zhu et al., 2026). We extracted hidden-state representations from a middle-to-late layer (layer 26) of GPT-J (Singh et al., 2023; Kantamneni & Tegmark, 2025; Stolfo et al., 2023) while the model processed the arithmetic stimuli, and we applied the same geometry analyses used above for the neural data (**Figure 3M-U**). Unlike in the human MTL, LLM representations of the numbers showed a curved, smooth, and magnitude-related progression (Kantamneni & Tegmark, 2025; Sun Y. et al., 2025; for a possible reason, see Karkada et al., 2026; **Figure 3M**). This numerical-distance structure was confirmed by RDM analysis (slope β = 2.4947; p[slope = 0] = 1.9e-46; **Figure 3P**). Importantly, however, a distance effect alone does not necessarily imply a one-dimensional number line: a representation can preserve increasing distances among numbers while remaining high-dimensional (see **Supplementary Figure 2** for an example). Consistent with this interpretation, transition-angle analysis showed that the LLM representation was much more folded than a noisy number-line control and closely matched the noisy-simplex control (**Figure 3N-O**). LLM embedding also had high shattering dimensionality (**Figure 3Q**).

To determine what drives the geometry of human MTL arithmetic representations, we evaluated two competing models at the single-subject level. For each subject, we correlated their trial to trial neural representational distance matrix (RDM) against both a scalar numerical-distance RDM and an LLM-derived RDM. A paired comparison of these fits revealed that the neural geometry aligned significantly better with the LLM RDM than with the numerical-distance RDM (numeric-distance correlation: −0.0099 ± 0.0087; LLM-RDM correlation: 0.0319 ± 0.006; two-sided paired test: p = 0.00198; **Figure 3R**). This result demonstrates that MTL arithmetic task representations are better described by a high-dimensional symbolic geometry, as instantiated by an LLM, than by simple scalar numerical distance.

The remaining LLM analyses paralleled the neural results. Cross-condition generalization across the 11 meaningful number dichotomies was limited, with most dichotomies generalizing in only 0–2 subjects or folds (mean CCGP: 0.5024 ± 0.0052; **Figure 3S**). Linear and log number-line projections explained substantially less trial variance than the residual high-dimensional condition-mean structure (linear number line: 8.4241 ± 1.1776%; residual condition mean: 33.6830 ± 2.1420%; p = 6.5e-10; log number line: 9.4102 ± 1.2913%; residual condition mean 32.6969 ± 2.0821%; p = 1.8e-09; **Figure 3T**). At the single-trial decoding level, the canonical number line was not the uniquely optimal ordering (best cross-validated linear regression decoding performance) among all possible number lines (relabel the 9 numbers in all possible 9! ways and repeat the decoder, **Methods**). The linear number-line rank was not significantly above chance (p = 0.0615) and was significantly below the best possible ordering (p = 4.88e-04), while the log number-line rank showed a modest above-chance bias (**Figure 3U**; p = 0.0049) but also remained far below the optimal ordering (p = 0.0020). Linear and log models did not differ significantly from each other (p = 0.0674). Thus, even in an artificial symbolic system that preserves stronger numerical-distance structure than the MTL (Hu et al., 2026), number representations remain high-dimensional and are only partially captured by a number-line projection.

### Implementation of arithmetic with simplicial numerical codes

How do simplicial representations of numbers afford arithmetic computations? We hypothesized simple arithmetic may function the way it does in large language models (LLMs): routing operand- and operator-related information via attention to the answer position, where a multi-layer perceptron (MLP) transforms this integrated information into result-related representations (Stolfo et al., 2023).

Under this general framework, we sought to determine the specific information integration strategy used to derive the result by comparing different model variants (**Figure 4A-F**). We define the neural result representation Y_res_ as the serialized population firing rate across multiple time bins during the thinking period (**Figure 4A**). Furthermore, we hypothesize that Y_res_ is linearly transformed from the two operand representations X_op1_ and X_op2_, which we define as the pooled temporal components from tuned neurons. Importantly, the transformation should happen in a context-specific (plus vs. minus) manner (**Figure 4B-C**). Indeed, it can be seen that in both the flexible (**Figure 4B**) and commutative (**Figure 4C**) model, X_op2_ are weighted differently depending on the sign of the operation. To test the possibility that the brain treats “3+4” equally as “4+3” by the commutative law, we shared weights between X_op1_ and X_op2_ during addition in the commutative model (**Figure 4C**). Two baseline models were developed where subjects simply memorized information from one of the operands, but incorrectly ignored information from the other operand, i.e., operand2 (**Figure 4D**) or operand1 (**Figure 4E**) to derive results. Finally, another control model simulated the case where the brain overlooks the mandate of specific arithmetic rules on how <u>operand1</u> and <u>operand2</u> should be integrated to derive result representations (**Figure 4F**).

**Figure 4.**
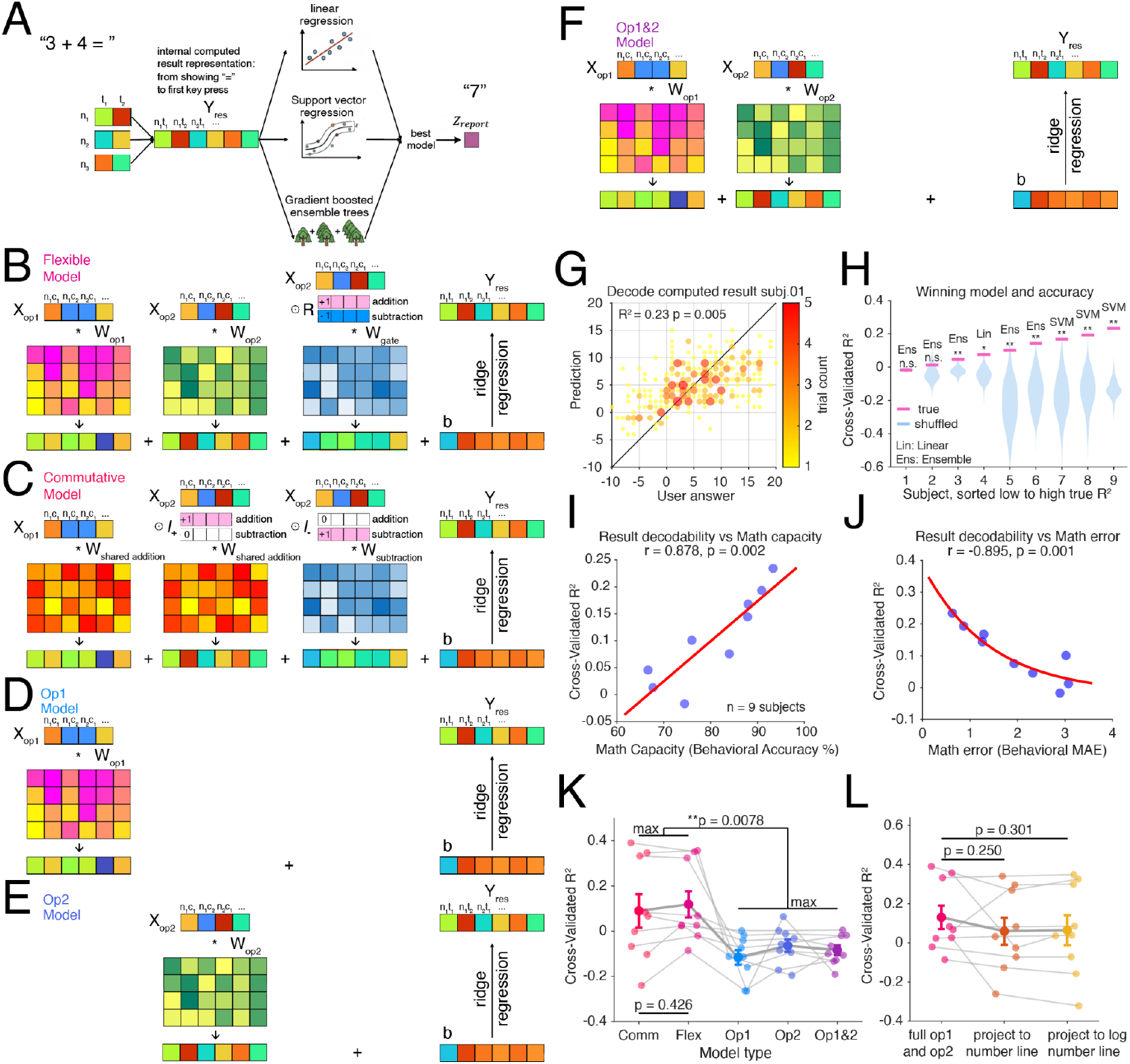
Implementation of arithmetic with simplicial numerical codes. **Abbreviations**: Comm, commutative model; Ens, ensemble decoder; Flex, flexible model; Lin, linear decoder; Op1, operand 1; Op2, operand 2; SVM, support-vector machine (regression) decoder; Yres, result-period neural representation. MAE, mean absolute error. W, weights of the regression model. **A**, Decoding framework for testing whether result-period activity encodes the internally computed answer. Population activity neuron-by-time bin spanning from the appearance of the equals sign to the first keypress is serialized into Yres. Linear, support-vector, and gradient-boosted ensemble tree regression decoders are trained to predict the participant’s reported answer. **B**, Flexible operation-dependent model. Yres is ridge regressed from operand representation Xop1 and Xop2 together with an operation-gated term such that operand information can be weighted differently for addition and subtraction. Xop1 and Xop2 are formed by temporal components *c* of the neurons *n* (serialized across all tuned neurons of a subject) **C**, Commutative model. Addition weights are shared across operands to induce commutativity, while an operation-specific interaction term allows subtraction-dependent transformations **D**, Operand-1 baseline model. The first-operand representation Xop1 is linearly transformed by a learned weight matrix and combined with a ridge-regression bias term to estimate the result-period neural representation Yres. **E**, Operand-2 baseline model. Same as panel D, but using only the second-operand representation Xop2. **F**, Operand-1-and-operand-2 baseline model. Xop1 and Xop2 are transformed independently and summed to estimate Yres, without explicitly incorporating the arithmetic operation (plus vs minus). **G**, Example subject showing decoded result value versus the participant’s entered answer (N = 300 trials). The diagonal indicates perfect prediction, and scatter color or size indicates trial count. **H**, Cross-validated result-decoding performance across subjects. Subjects are sorted by true decoding performance; pink markers indicate true-label performance, blue violins indicate label-shuffled null distributions, and text labels denote the winning decoder class. N = 9 subjects that performed both addition and subtraction. N = 200 shuffles. Negative R-squared means that the prediction is worse than guessing the mean. **I**, Across-subject relationship between result decodability in H and behavioral math capacity. Each point represents one subject, and the red line shows the fitted linear trend. Pearson correlation r and corresponding p-value were reported. **J**, Across-subject relationship between result decodability and behavioral math error (mean absolute error between ground truth to reported result). Each point represents one subject, and the red curve shows the fitted exponential trend. Pearson correlation r and corresponding p-value were reported. **K**, Comparison of candidate arithmetic-transformation models in B-F. gray lines connect the same subjects, and error bars indicate mean ± SEM. The best operation-dependent model (Flex and Comm) outperforms the best baseline models, whereas the commutative and flexible models are not clearly distinguishable, color indicates different models. **L**, Effect of projecting operand representations Xop1 and Xop2 onto one-dimensional number-line axes before estimating Yres. Performance for the full (B-F) operand representation is compared with models using linear and logarithmic number-line projections; gray lines connect subjects, and error bars indicate mean ± SEM. Number-line projection does not improve the transformation from operand representations to decodable result representations.

To understand whether thinking period population neural activity in MTL actually captures upcoming internal computed results, we fit decoder models of different complexities to read out upcoming answers the subjects entered (**Figure 4A**). Indeed, MTL population dynamics can be used to decode mentally computed results, where the decoded answer largely matches the subject’s answer in one example (**Figure 4G**). Interestingly, across all 9 subjects that performed both addition and subtraction, the best model was nonlinear in 8/9 cases (**Figure 4H**), suggesting that nonlinear transformations are likely present to convert MTL activity to final behavioral readout. Importantly, the degree to which we can read out mentally computed results from MTL positively correlated with the behavioral accuracy of different subjects (p < 0.01 **Figure 4I**) and negatively correlated with the average magnitude of the error (p < 0.01 **Figure 4J**) when incorrect answers were given. This might be indicative that when MTL activity fails to meaningfully decode mentally computed results against a label shuffled control in 2/9 subjects, it’s not because that MTL activity lacks result representation in general, but more likely that the subjects were generally unsure what to answer, potentially adjudicating between multiple answers, which also confused the decoders. Finally, when the estimated Y_res_ (**Figure 4A-E)** was fed into the decoder, we found insufficient evidence that either the flexible or commutative model was the preferred strategy the brain uses for the arithmetic transforms (p = 0.43 Wilcoxon signed-rank test **Figure 4K**). Yet the better of the two models (flexible and commutative) clearly outperformed the best of the baseline, ablated models (p < 0.001, Wilcoxon sign rank test **Figure 4K**), indicating that the result representation is indeed utilizing both operands and the operation information. One important property of simplex-like representations is that they can be projected into 1-d line-like representations of arbitrary order; it’s intuitive to hypothesize that the particular arithmetic task can be solved with a mental number line-like representation. Therefore, we explore whether projecting X_op1_ and X_op2_ onto a 1-d number line representation through linear regression before transforming them into mentally derived result representation is a better strategy for mental arithmetic. Interestingly, projecting to neither the linear nor the log number line yielded significant improvement (**Figure 4L**, p < 0.30, two-sided Wilcoxon sign rank test) in transforming operand representation into mentally computed result representation that is predictive of what the subjects eventually entered.

## DISCUSSION

We find that coding for numbers 1-9 has a simplicial manifold structure in the human MTL. Whereas much of human neuroscience has focused on linear magnitude as the organizing principle for numbers, the neural geometry we observe is better aligned with the idea that mentally, numbers are organized along multiple dimensions (Shepard et al., 1975). Relative to a low-dimensional manifold, like a number line, a simplex offers several advantages. First, it has **high shattering dimensionality**, which permits zero-shot discrimination for any numerical subset (e.g., even/odd, prime/composite). Second, it has high **expressivity**, which permits zero-shot reordering of the digits. This ordering includes the standard 1-9 ordering; consequently, a simplicial manifold is a more general and more flexible representational scheme than a number line. Third, in the case of a regular simplex, its equidistance makes it more robust to noise and interference. A rich and influential body of literature has firmly established the existence of a mental number line (Dehaene et al., 1999; Dehaene et al., 2008; Knops et al., 2009; Rugani et al., 2015), including at the level of single neurons in the MTL (Kutter et al., 2018, 2024). Our result extends this perspective to high dimensionality.

In both artificial and biological networks, learning reshapes representational geometry by expanding the global dimensionality of the latent space to decorrelate heavily trained concepts (Flesch et al., 2022; Sun W. et al., 2025; Nguyen et al., 2024; Failor et al., 2025). From early childhood, individuals are exposed to both counting and Arabic numbers and a broad range of numerical manipulations (especially 1–9, Barner, 2017; Slusser, Santiago, & Barth, 2013; Carey, 2009; Gallistel & Gelman, 1992; Spelke, 2017). This intensive, lifelong overlearning of small numbers is therefore expected to drive the underlying neural manifolds to expand into a maximal-dimensional configuration (Nakai, et al., 2023; Chung et al., 2018). This idea is also consistent with our finding that LLM representations of numbers also have an inherently high dimensional structure. By directly quantifying this dimensionality, our results further extend prior LLM work that describes the geometry of numbers only in dimension-reduced spaces (Kantamneni & Tegmark, 2025; Sun Y. et al., 2025; Karkada et al., 2026; Hu et al., 2026).

Whether symbolic and nonsymbolic numbers share a common representation remains debated. Evidence for partially distinct systems comes from developmental and comparative work showing approximate numerosity representations appear before, or outside, formal symbolic math training (Starr et al., 2013; Nieder et al., 2002; Nieder et al., 2006; Rugani et al., 2015), whereas exact symbolic representations suitable for arithmetic appear later in life and are highly influenced by cultural, educational factors (Ferrigno et al., 2017; Pica et al., 2004; Dehaene et al., 2008; Dillon et al., 2017; Banerjee et al., 2025). However, behavioral links between approximate quantity and formal mathematics suggest that symbolic and nonsymbolic systems could have shared representation or are at least partially coupled (Gilmore et al., 2007; Halberda et al., 2008; Ferrigno et al., 2017). The neural basis of this relationship remains equally debated. Some work supports shared magnitude representations across nonsymbolic and symbolic formats, especially in intraparietal cortex, whereas other studies reveal notation-, hemisphere-, region-, and task-dependent numerical codes (Ansari, 2016; Aulet, Kaicher, & Cantlon, 2025; Cantlon et al., 2009; Kersey, Aulet, & Cantlon, 2026; Piazza et al., 2007; Cohen Kadosh et al., 2007; Bulthé et al., 2014; Sokolowski et al., 2017; Wilkey et al., 2020; Diester & Nieder, 2007; Seidler et al., 2026; Emerson & Cantlon, 2015). In human MTL, prior work established that nonsymbolic and symbolic numbers are represented differently by single neurons (Kutter et al., 2018). We similarly found that number codes were format dependent. Critically, however, we provide a new perspective to this question by showing that despite the apparent use of different codes, the nonsymbolic and symbolic number representations share a partially aligned latent representational subspace identifiable by SVCCA.

This provides a potential computational mechanism that allows us to flexibly transfer between the nonsymbolic and symbolic numbers, e.g., buying 5 apples (non-symbolic quantities) with a bill that reads $5 (symbolic). Here we tested linear alignment; future work with larger within-task trial budgets and better matched tasks between formats could test whether nonlinear transformations support shared neural representation between formats (Andrew et al., 2013; Radford et al., 2021).

How the brain transforms symbolic operands into arithmetic results remains unresolved (Nieder, 2025; Qin et al., 2014; Pinheiro-Chagas et al., 2024), although one study did identify abstract operation rule-selective neurons in MTL (Kutter et al., 2022). One possible reason such calculation result information may have been missed is methodological choice: many prior analyses pooled neurons/channels across patients into pseudo-populations. This approach is well suited for detecting general coding principles, but it disrupts the link between an individual subject’s neural state and that subject’s own reported answer. This distinction is important for arithmetic, where subjects differ in strategy and math capacity. Indeed, we found that subjects with weaker result decodability also showed lower behavioral accuracy and larger errors. Motivated by recent work on arithmetic in large language models, we separated the computation into two stages. In transformer models, attention can route operand- and operator-related information to the answer position, while MLP modules transform that merged information into result-related representations (Stolfo et al., 2023). By analogy, we asked whether MTL activity during the thinking period reflects an operation-dependent transformation of operand representations, and whether this activity can then be decoded into the answer that subjects eventually report.

Our findings support this computational decomposition. Analogous to an attention mechanism integrating contextual information, MTL activity cannot be explained by the mere working memory traces of isolated operands; it requires their operation-specific integration. Similar to an MLP module transforming these inputs into an output, MTL population activity can nonlinearly decode arithmetic results (which has never been shown to our knowledge), with decoding strength directly predicting the subject’s behavioral accuracy. These findings extend recent fMRI evidence showing neural encoding of internally produced quantities (Czajko et al., 2024). Crucially, our work advances this principle from approximate, single-operand transformations of nonsymbolic quantities to exact, multi-operand symbolic arithmetic.

What neuron-level tuning functions produce simplicial manifolds? One possibility is a labeled-line (i.e., one-hot) code, in which each numerosity is represented by neurons selective only for that number. However, another possibility is that individual neurons do not systematically encode specific numbers; indeed, they could have complex multi-peaked tuning curves (Foucault et al., 2025; Mainali et al., 2025). Such tuning curves are typically diagnostic of complex, warped, or folded manifold structures, and are often associated with function superposition, in which multiple dissimilar, non-adjacent concepts are overloaded onto single neurons (Elhage et al., 2022; Maimon et al., 2026; Yan et al., 2026b).

Inspection of our single-neuron tuning curves argues strongly against the preferred-number regime and in favor of the alternative, suggesting a mechanism by which the brain maximizes numerical capacity with a finite number of neurons.

Future studies will be needed to address the limitations of the present work. First, and most important, a large body of work demonstrates the central role of the parietal lobe and lateral frontal cortex for numerical cognition (Nieder & Dehaene, 2009). It remains an open question whether the simplicial manifold structure we identify here will extend to these regions. Second, it remains unclear whether the principles we identify for 1-9 will apply to larger numbers. We deliberately restricted our stimulus set to single-digit non-zero numbers, so any suggestions about how the brain represents other higher or lower numbers are necessarily speculative. In any case, it seems likely that the brain does not simply increase dimensionality infinitely for all higher numbers. Third, the neural processing for more complex arithmetic rules like multiplication or division remains an exciting unknown avenue. Finally, since counting small quantities and simple arithmetic are more heavily practiced in adults, future developmental work is needed to trace the trajectory of this simplicial structure throughout the gradual acquisition of mathematical skills (Kersey & Cantlon, 2017; Kersey, Aulet, & Cantlon, 2026; Menon, 2010).

## Acknowledgements

We thank Xue-Xin Wei, Robbe L. T. Goris, Joshua Adkinson, Victoria Gates, George Kokalas, Raissa Mathura, Layth Mattar, and Andrew Watrous for their assistance and advice.

## Supplementary Figures

**Supplementary Figure 1.**
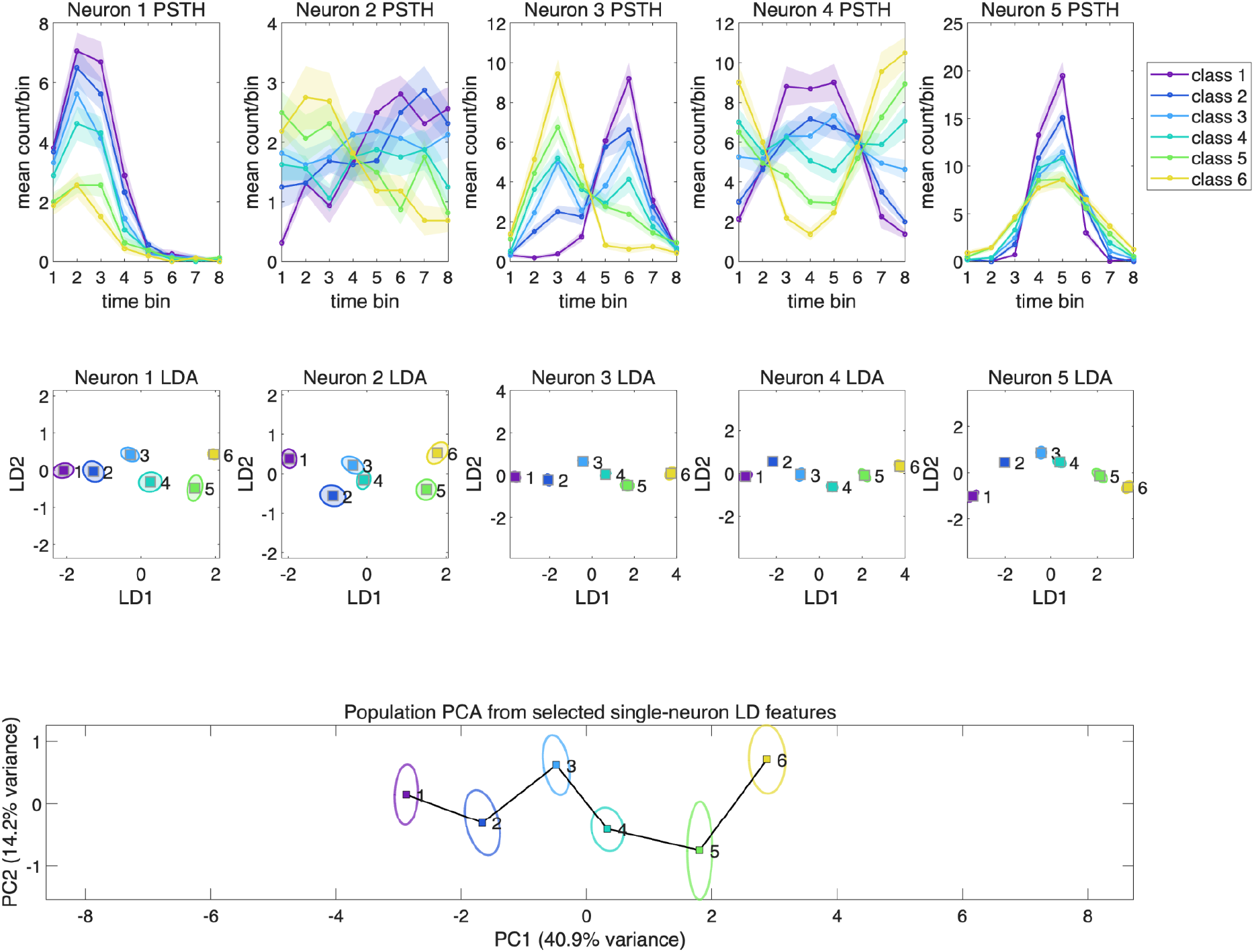
LDA-based temporal component extraction does not by itself create a population structure that strongly deviates from a number-line. Top row, mean PSTHs for five **simulated** neurons across ordered numerical classes; shaded regions indicate SEM across trials. Middle row, the corresponding single-neuron LDA projections computed from each neuron’s trial-by-time-bin response, with ellipses indicating SEM around each class mean and square markers indicating class centers, as in Figure 1 H,I,L. Bottom row, PCA of the five LD1/LD2 features from the five single neurons after z-scoring, as in Figure 2 D,N, but in 2D for simplicity. The connected class centers form an ordered, approximately one-dimensional number-line-like trajectory, with the variance explained by PC1 and by PC1+PC2 shown on the plot. This simulation shows that when neurons contain condition-dependent temporal dynamics that vary smoothly with numerical class, LDA-based temporal component extraction can recover the underlying ordered structure rather than artificially producing a higher-dimensional or non-number-line manifold. Thus, the use of LDA for temporal feature extraction cannot by itself be interpreted as necessarily inflating dimensionality.

**Supplementary Figure 2.**
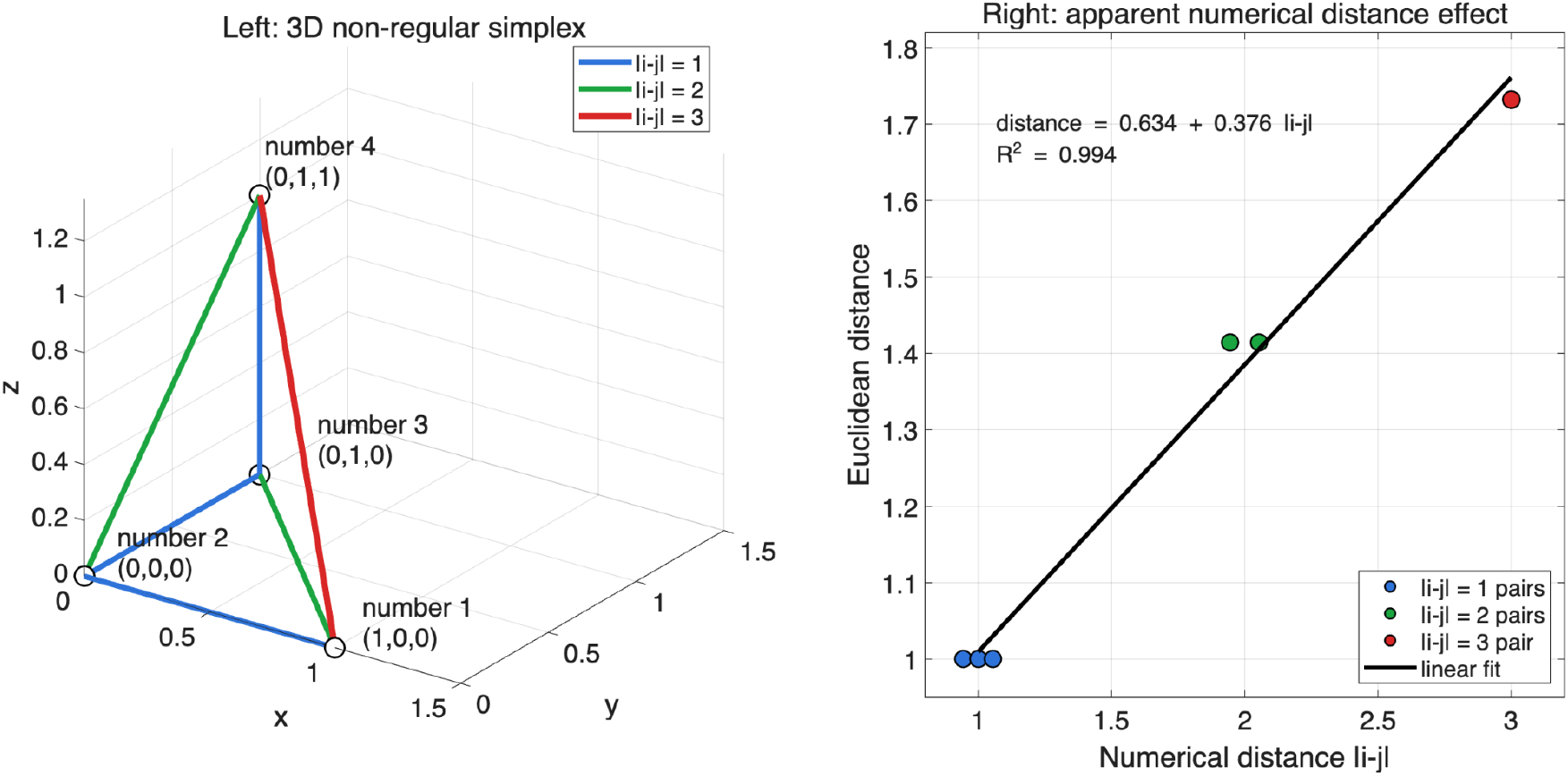
A numerical distance effect does not imply a line-like number representation. Left, a fixed non-regular 3-simplex with vertices (0,0,0), (1,0,0), (0,1,0), and (0,1,1), assigned to numbers 1–4. Edges are colored by numerical distance |i−j|, and edge annotations show Euclidean distances between vertices. Right, Euclidean distance between pairs of vertices plotted against numerical distance |i−j|, with a linear fit. The structure is clearly three-dimensional, not a line, yet it still produces an apparent numerical distance effect. Thus, a numerical distance effect alone is not sufficient evidence for a line-like number representation.

**Supplementary Figure 3.**
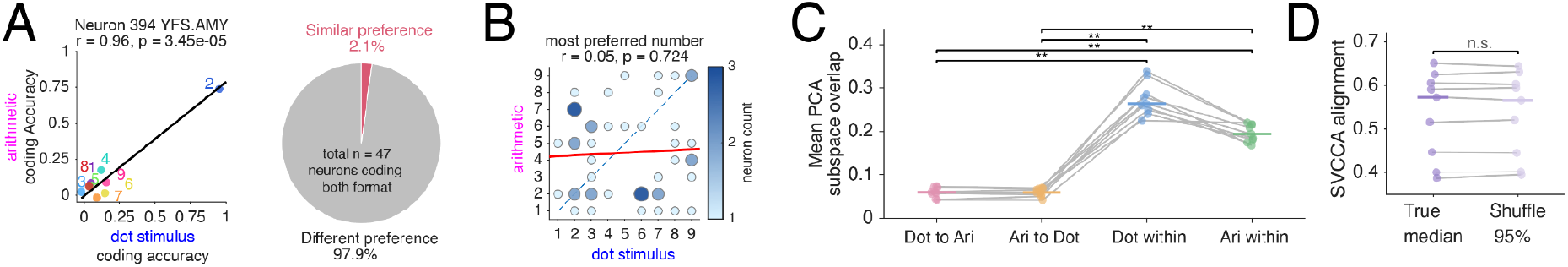
Number codes from dot-counting and arithmetic tasks are distinct and not reliably linearly aligned. **A**, Left: Example neuron coding both dot-stimulus numerosity during the counting task and number information during the arithmetic task. Each point shows coding accuracy for one number in the two conditions; colors indicate number identity, and the black line shows the best linear fit. Right: Fraction of neurons coding both dot-counting and arithmetic-task number information that showed the same preferred number across conditions. Only a small fraction of dual-coding neurons showed matched number preference, indicating that single-neuron selectivity was usually different across the two task/format contexts. **B**, Preferred-number matrix for neurons coding both dot-stimulus numerosity and arithmetic-task number information. Circle size and shading indicate neuron count, the dashed diagonal marks identical preferred numbers across conditions, and the red line shows the linear trend. Preferred numbers were not correlated across the two representations. **C**, Population subspace overlap between dot-counting and arithmetic number codes. Cross-condition overlap was measured in both directions and compared with split-half within-condition overlap for each representation. Gray lines indicate subjects, colored markers show subject-level values or means, and brackets indicate paired statistical comparisons. Cross-condition overlap was significantly lower than within-condition overlap, showing that the two representations occupied largely distinct population subspaces. **D**, SVCCA alignment between dot-counting and arithmetic population codes compared with a label-shuffled control. Gray lines indicate subjects, and points show the true median alignment and the shuffled-control 95th percentile. The true alignment did not significantly exceed the shuffled control, indicating that number codes expressed in different formats and different tasks were not reliably linearly aligned. **P < 0.01; n.s., not significant.

## Methods

### Human intracranial neurophysiology

Intracranial electrophysiological recordings were obtained from adult patients undergoing clinical intracranial monitoring for medically refractory epilepsy at Baylor St. Luke Hospital, following established intracranial recording procedures during epilepsy monitoring (Xiao et al., 2024; Franch et al., 2026a). Electrode placement was determined exclusively by clinical criteria. All participants provided informed consent under protocols approved by the Baylor College of Medicine institutional review board.

Single-neuron activity was recorded from stereotactic depth electrodes equipped for unit recordings, including AdTech Medical Behnke–Fried–style probes with 8 channels per probe. Electrode locations were verified by co-registering each participant’s pre-operative anatomical MRI with the post-operative CT scan (Franch et al., 2026a).

Each depth probe contained eight microelectrode channels optimized for unit recording. Neural signals were acquired using a 512-channel Blackrock Microsystems NeuroPort system at 30 kHz. Spikes were detected and sorted offline with Wave_clus (Chaure et al., 2018), followed by manual curation. Putative units were retained when their waveforms were stable over the recording session and showed consistent morphology, including amplitude and trough-to-peak features, together with inter-spike-interval distributions consistent with a refractory period. Units with inter-spike intervals violation of more than three percent were defined as multi-unit. Unless otherwise stated, analyses included both single-unit and multi-unit activity (Franch et al., 2026a).

### Electrode localization

Electrode localization was performed using a standardized MRI/CT co-registration workflow (Franch et al., 2026a). For each patient, pre-operative T1-weighted MRI and post-operative CT images were converted to NIfTI format (Li et al., 2016). The post-operative CT was aligned to the pre-operative MRI using FSL (Jenkinson & Smith, 2001; Jenkinson et al., 2002). Electrode contacts were then manually identified in BioImage Suite (Joshi et al., 2011) and transformed into each participant’s native anatomical space using iELVis MATLAB utilities (Groppe et al., 2017; Yang et al., 2012).

Cortical surfaces were reconstructed using FreeSurfer (Dale et al., 1999), and electrode contacts were visualized on each participant’s native anatomy using iELVis (Groppe et al., 2017). Microelectrode positions were defined relative to the deepest macro-contact of the Behnke–Fried depth electrode assembly. For group-level visualization, participant anatomies and electrode coordinates were warped into MNI152 space using RAVE (Magnotti et al., 2020).

### Behavioral tasks

Participants performed visually presented numerical tasks during intracranial recording. Stimuli were generated and presented using MATLAB with Psychtoolbox on the center spherical region of a 124 × 71.5 cm (W x H) TV with a refresh rate of 60 Hz. Participants viewed the screen from approximately 170 cm away. Stimuli were presented on a black background, and participants responded using a standard numeric keypad with numbers arranged in four rows by three columns. The keypad contained additional response keys (e.g., enter/delete).

Stimulus timing was synchronized with the electrophysiological recordings using a photodiode taped to the corner of the display. A small luminance patch in that corner changed intensity at task-event onsets, and the photodiode signal was recorded together with the neural data to align behavioral events with electrophysiological recordings.

### Dot-counting task

In the dot-counting task, participants viewed arrays of white dots and reported the number of dots using the numeric keypad. Dot diameter was 1.5cm on screen, which is about 0.51 degrees of visual angle (similar to the visual size of a full moon on sky), and dot locations were randomized independently on each trial. Dots were prevented from having spatial overlap with other dots.

Each trial began with presentation of the dot array for 1250 ms, followed by a blank delay period of 500 ms. A response prompt was then displayed, and participants entered their response at their own pace. After the response, visual feedback of entered answer was shown for 500 ms (in red when the answer is wrong and in green when the answer is correct), followed by a 500 ms inter-trial interval before the next stimulus. The position of the dots was randomized.

In one parameter set, 4 patients completed trials using numerosities 1–10, with each patient contributing 30.00 ± 0.00 trials per number on average. In a second parameter set, 6 patients completed trials using numerosities 1–16, with each patient contributing 8.63 ± 0.95 trials per number on average. All numbers in mean + s.e.m.

### Arithmetic task

In the arithmetic task, participants solved visually presented addition and subtraction problems. Each trial contained two operands and an operation sign, presented sequentially. The size of a single digit numeral is about 3 cm by 5 cm (W X H). The three task items were each displayed for 750 ms. The operation sign could appear in the first, second, or third serial position yet the effect of the position is investigated in a separate study and beyond the scope of the current work. Regardless of the position of the operation sign, participants were instructed (during subtraction trials) to subtract the second shown number from the first shown number, i.e., to compute first number − second number.

After the final item in the sequence, a response prompt was shown, and participants entered their answer at their own pace using the numeric keypad. Visual feedback was presented for 500 ms of the entered number being green (correct) or red (incorrect), followed by a 500 ms inter-trial interval.

For the arithmetic task, in one parameter set, 4 patients completed trials using numbers 0–10, with each patient contributing 41.82 ± 7.35 trials per number on average. In a second parameter set, 7 patients completed trials using numbers 0–16, with each patient contributing 12.45 ± 0.69 trials per number on average. All numbers in mean + s.e.m.

### Single-neuron firing-rate and temporal-component decoding during dot counting task

Single-neuron number decoding was performed separately for the arithmetic and dot-counting tasks. For dot counting task, spike trains were aligned to stimulus onset (i.e. dot array), and all decoding analyses used spikes occurring from 0.1 to 1.9 s after the alignment event. This yielded a 1.8-s analysis window. Trials were assigned to number conditions using the ground truth number of dots for each trial.

For each subject, number labels were sorted in ascending order, and the first nine number conditions with at least two trials were selected (for all other than 1 subject, this results in 1-9, where the remaining subject included 10 and 11 but excluded 2 and 5). Trials whose labels fell outside this selected set were excluded from analysis. The analyses reported here were therefore nine-way number decoding analyses, with a chance level of 1/9.

### Firing-rate representation

For the firing-rate code, each trial from a given neuron was represented by a single value: the total spike count in the 0.1–1.9 s analysis window. For plotting firing-rate tuning curves, this spike count was converted to firing rate by dividing by the 1.8-s window duration. This representation therefore discarded within-window temporal structure and retained only the overall response magnitude for each trial.

### Temporal-window representation

For the temporal code, the same 0.1–1.9 s analysis window was divided into non-overlapping time bins, and spikes were summed within each bin. Each trial was therefore represented as a vector of spike counts across time for one neuron.

We scanned candidate bin lengths of 90, 120, 180, 225, 300, 360, 450, 600, 900, and 1800 ms. Because the full analysis window was 1.8 s, these bin lengths corresponded to 20, 15, 10, 8, 6, 5, 4, 3, 2, and 1 temporal bin(s), respectively. The 1800-ms condition was included as the one-bin, whole-window case, allowing the temporal code representation to include the firing-rate representation as a special case (limit).

### LDA temporal components

For each neuron, number-discriminating temporal components were estimated using multiclass linear discriminant analysis. Let

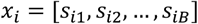

denote the temporal response vector for trial *i*, where *s*_*ib*_ is the spike count in temporal bin *b*, and *B* is the number of bins for a given candidate bin length. LDA estimates linear combinations of these temporal bins that maximize separation between number conditions relative to within-condition variability. Each temporal component therefore corresponds to a weighted temporal pattern over the analysis window, and each trial can be projected onto these components.

For nine-way decoding, LDA can produce up to min (*B*, 8) discriminant dimensions, where *B* is the number of temporal bins and 8 is the maximum number of discriminant dimensions for nine classes. For visualization, trials were projected onto the first two temporal components when at least two components were available. These projections were used to visualize temporal separation among number conditions at the single-neuron level. Quantitative decoding performance was always estimated using held-out trials in cross-validation without sub-selecting the number of components to use.

### Cross-validated decoding

For each neuron, we trained a regularized linear discriminant classifier to predict the number condition from either the firing-rate representation or the temporal-window representation, similar to those used in (Zhu et al., 2025). The classifier (matlab’s fitcdiscr) used a linear discriminant model with a uniform class prior, so that unequal trial counts across number conditions did not bias the classifier toward more frequently presented numbers. We scanned the temporal bin lengths listed above together with LDA shrinkage parameters Gamma:

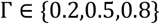

For each neuron, the bin length and Γ value that jointly produced the highest cross-validated decoding accuracy were selected as that neuron’s temporal-code parameters. Features with zero within-class variance in the training data were removed before fitting the classifier.

Decoding accuracy was estimated using stratified cross-validation. We requested 10-fold cross-validation. When the trial counts for a session did not permit 10 folds while preserving all number classes in each test fold, the number of folds was reduced to the largest value that maintained at least one trial from every number condition in each fold. On each fold, the classifier was trained on the training trials and evaluated on the held-out trials. Overall decoding accuracy for a neuron was defined as the fraction of held-out trials whose predicted number matched the true number:

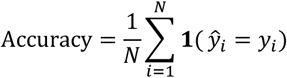

where *y*_*i*_ is the true number label for trial *i*, ŷ_*i*_ is the predicted number label, and *N* is the total number of held-out predictions accumulated across cross-validation folds.

### Number-specific decoding accuracy

In addition to overall decoding accuracy, we computed number-specific decoding accuracy for each neuron. After cross-validation, the predicted labels from held-out trials were placed back into their original trial order. For each number *c*, per-number accuracy was computed as the fraction of trials whose true label was *c* and whose predicted label was also *c*:

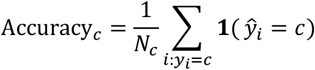

where *Nc* is the number of trials with true label *c*. This procedure yielded, for every neuron, both an overall cross-validated decoding accuracy and a separate cross-validated decoding accuracy for each number condition.

### Definition of number-coding neurons

Statistical significance of number decoding was assessed separately for each neuron using a label-shuffle permutation test. After the observed decoding accuracy was computed, number labels were randomly permuted across trials while preserving the neural data and the total number of trials assigned to each label. Cross-validated decoding was then repeated using these shuffled labels.

For the temporal-code analysis, the permutation test used the temporal bin length and Γ value selected from the observed data for that neuron. Thus, the shuffle distribution tested whether the selected temporal representation carried more number information than expected from random assignment of number labels to trials. For the firing-rate analysis, the same one-feature firing-rate representation was used for both the observed and shuffled analyses.

We used 200 label shuffles for each neuron. A one-sided permutation p-value was computed as:

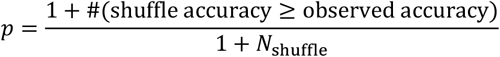

A neuron was classified as number-coding if its observed cross-validated decoding accuracy exceeded the shuffle distribution at *p* < 0.05

### Regional distribution of number-coding neurons

To test whether number-coding neurons were preferentially concentrated in particular medial temporal lobe regions, we compared the prevalence of temporal-component number tuning across regions. This analysis was performed separately for the arithmetic and dot-counting tasks.The previous section defined for each neuron:

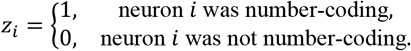

For visualization, we first computed the percentage of number-coding neurons separately for each subject and region:

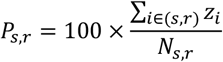

where *N*_*s,r*_ is the total number of recorded neurons from subject *s* in region *r*, and the numerator is the number of those neurons that met the temporal-component tuning criterion. Subject–region combinations with no recorded neurons were treated as missing values and were not included in the corresponding regional percentage summary.

For statistical testing, we did not fit the model to the subject-level percentages. Instead, we used a neuron-level binomial generalized linear mixed-effects model, which preserves the binary outcome for each neuron while accounting for repeated sampling of neurons within the same subject. The model was:

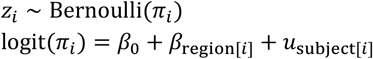

where *β*_region[*i*]_ is the fixed effect of anatomical region, and *u*_subiect[*i*]_ is a random intercept for subject:

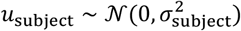

In MATLAB notation, the model was:

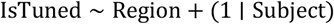

with a binomial distribution and logit link using the function fitglme. This model tested whether the probability that a neuron was classified as number-coding differed across regions, while accounting for the fact that neurons from the same patient are not independent. The main effect of region was assessed using an ANOVA on the fitted generalized linear mixed-effects model. Separate models were fit for the arithmetic and dot-counting tasks.

### Three-dimensional visualization of population geometry

For the example-subject visualization in Figure 2D, we constructed a population activity matrix from the temporal-component representation described above. For each neuron, trials were projected onto the neuron’s optimized temporal LDA components. Neurons were included if they passed the lenient tuning threshold used for population geometry analyses: *p* < 0.125. The resulting trial-by-feature matrix was formed by concatenating the retained temporal-component projections across tuned neurons:

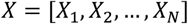

where *X*_*C*_ denotes the retained temporal-component (max 3 per neuron) from neuron *n*, and *N* is the number of included number-selective neurons. Principal-component analysis was then applied to the z-scored trial-by-feature matrix, and each trial was projected onto the first three principal components:

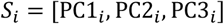

These three-dimensional PC scores were used only for visualization.

For each number condition *c*, we collected all trials with label *c* and computed the mean location in the three-dimensional PC space:

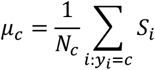

where *N*_*c*_ is the number of trials with dot number *c*. We also computed the within-condition covariance matrix:

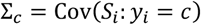

Each number-specific activity distribution was displayed as an ellipsoid centered at *μ*_*c*_. The ellipsoid was generated by eigendecomposing the covariance matrix,

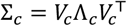

and transforming a unit sphere according to:

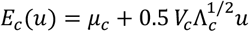

where *u* is a point on the unit sphere. Thus, the ellipsoid axes were aligned with the principal directions of the number-specific covariance, and their radii were scaled to 0.5 times the corresponding within-condition standard deviations. The ellipsoids were plotted as wireframe surfaces and colored by dot-number condition.

This visualization was descriptive and was not used for statistical inference. Its purpose was to show the geometry of the number-conditioned population activity in a low-dimensional PC projection and to illustrate whether number-condition centroids appeared to follow number line-like trajectory.

### Arithmetic operand-aligned representations

For analyses of number representations during the arithmetic task, neural activity was aligned to the onset of the numerical operands rather than the exact cue order shown on the screen. This is because the operation sign could appear in different orders of the three cues needed for the arithmetic, we first converted screen cue times into conceptual operand times.

When the operation sign appeared first, operand 1 occurred at the second screen cue and operand 2 occurred at the third screen cue. When the operation sign appeared in the middle, operand 1 occurred at the first screen cue and operand 2 occurred at the third screen cue. When the operation sign appeared last, operand 1 occurred at the first screen cue and operand 2 occurred at the second screen cue. Thus,

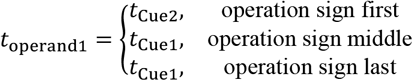

and

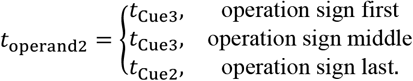

The operand-1-aligned and operand-2-aligned data were then concatenated along the trial dimension, so that each original arithmetic trial contributed two operand presentations. After concatenation, the analysis in Figures 1-3 ignored whether a number appeared as operand 1 or operand 2. Each operand presentation was labeled only by the corresponding numerical value.

### Stimulus-defined numeral representation for Figures 1–3

For Figures 1–3, we used a stimulus-defined numeral representation. In this analysis, operand labels were determined by the displayed numeral stimuli and were not adjusted based on the participant’s behavioral response. Thus, if a participant responded incorrectly, the trial was still labeled according to the numbers that were shown on the screen. For each subject, non-zero numerals 1–9 was selected, giving a nine-way decoding analysis with chance accuracy equal to 1/9. Trials with operand values outside the selected class set were excluded from analysis for Figures 1–3.

### Response-adjusted operand labels for Figure 4

For Figure 4, we constructed a response-adjusted operand representation for error trials intended to approximate the numerical structure used by the participant on each trial. This analysis was motivated by the fact that some arithmetic errors appeared consistently within and across subjects in a predictable way, such as using the wrong operation sign, potentially due to the short presentation period and randomized location of the stimuli, leaving subject without enough time to locate and recognize the task stimuli. Three response-adjustment rules were used.

First, if the response was consistent with using the opposite operation sign, the operation sign for that trial was switched. Operand order was not changed under this rule. For example, if the displayed operands were 5 and 3 on a subtraction trial, the task-defined answer was 5 − 3 = 2. If the participant responded 8, this response was consistent with computing 5 + 3 rather than 5 − 3, so the trial was treated as an operation-sign error and the operation label was switched from subtraction to addition. Similarly, a response of 6, 7, 8, 9, or 10 would fall within the ±2 response-unit tolerance around the alternative answer 8.

Second, when the minus sign appeared before the operands and the response was consistent with attaching the minus sign to the first operand, the first operand was relabeled as a signed negative value. In this case, the expression was represented as −operand1 + operand2. For example, if the display sequence was “− 5 3,” the task-defined answer was 5 − 3 = 2. If the participant responded −2, this response was consistent with treating the expression as −5 + 3. For the response-adjusted representation, the first operand was therefore relabeled from 5 to −5, the second operand remained 3, and the operation was represented as addition.

Third, when the minus sign appeared after the operands and the response was consistent with computing operand 2 minus operand 1, the operand order was treated as reversed. For example, if the display sequence was “5 3 −,” the task-defined answer was 5 − 3 = 2. If the participant responded −2, this response was consistent with computing 3 − 5. For these trials, the neural snippet aligned to the displayed first operand, 5, was treated as the operand-2 response, and the neural snippet aligned to the displayed second operand, 3, was treated as the operand-1 response. The corresponding operand labels were swapped before subsequent pooling or projection.

For each candidate error type, the participant’s response was compared with the answer predicted by that alternative parse. A trial was assigned to an error type when the response was within 2 response units of the predicted alternative answer. When more than one alternative parse matched the response, the parse with the smallest absolute deviation from the participant’s response was selected. For example, if the display sequence was “− 1 6,” the task-defined answer was 1 − 6 = −5. The opposite-sign parse predicted 1 + 6 = 7, whereas the minus-attached-to-first-operand parse predicted −1 + 6 = 5. If the participant responded 5, the response was within 2 units of both alternatives, because |5 − 7| = 2 and |5 − 5| = 0. Because the deviation was smaller for the minus-attached parse, the trial was assigned to the second error type.

Remaining exact ties were resolved using the participant’s dominant error pattern. For example, in an ambiguous case such as “− 2 4,” the task-defined answer was 2 − 4 = −2, the opposite-sign parse predicted 2 + 4 = 6, and the minus-attached parse predicted −2 + 4 = 2. A response of 4 is exactly 2 units away from both alternative parses. In such cases, the trial was assigned according to the participant’s dominant pattern among non-tied error trials.

Finally, error trials in which the participant’s response appeared to be simple copying of a visually presented number were not used as evidence for label adjustment and thus not adjusted. For example, if the display sequence was “7 2 −,” the task-defined answer was 7 − 2 = 5. A response of 7 would copy the first displayed operand. Even though this response is within 2 units of the wrong-sign answer 7 + 2 = 9, it was treated as a copying-like error rather than evidence that the participant systematically used the opposite operation sign.

After these response-based adjustments, operand labels were defined from the adjusted trial label than from the originally displayed digit labels alone. Therefore, the Figure 4 representation was not restricted to 1–9. Candidate values for LDA fitting were selected from the adjusted operand labels. Values were included in the fitting set only if they occurred twice. Zero was also included as a valid class for Figure 4.

### Temporal features for arithmetic operands

Because individual arithmetic cues were displayed for a shorter duration than the dot arrays, arithmetic operand decoding used a shorter post-onset analysis window. For each operand presentation, decoding features were extracted from 0.05 to 0.95 s after operand onset, yielding a 0.9-s analysis window.

For the firing-rate representation, each operand presentation for a given neuron was represented by the total spike count in this 0.05–0.95 s window.

For the temporal representation, the same 0.05–0.95 s window was divided into non-overlapping temporal bins. Spikes were summed within each bin, so that each operand presentation was represented as a vector of spike counts across time. We scanned candidate bin lengths of 60, 75, 90, 100, 150, 180, 225, 300, 450, and 900 ms. Because the full analysis window was 0.9 s, these bin lengths corresponded to 15, 12, 10, 9, 6, 5, 4, 3, 2, and 1 temporal bin(s), respectively.

The 900-ms condition was included as the one-bin whole-window case. Thus, the temporal analysis included the firing-rate-like limit and was not forced to use a multi-bin representation. The LDA fitting, temporal projection, cross-validation, and significance procedures were otherwise consistent with those used in the dot-counting task. (described above).

### Projection of additional adjusted values

For Figure 4, LDA fitting was restricted to operand values with sufficient repetitions, defined as at least two presentations after response-based adjustment. Operand values with insufficient repetitions were not used to optimize temporal bin length, shrinkage, or decoding accuracy. After each neuron’s optimized LDA model was fit using the eligible operand values, remaining operand presentations were projected through the learned temporal components for downstream representational analyses.

### Transition-angle analysis of number geometry

To quantify whether number representations were more consistent with an ordered number line or a simplex-like geometry, we measured transition angles between sequential number conditions. Analyses were performed separately for each subject using the same population representation used for the Figure 2 geometry analyses. Unless otherwise stated, analyses were restricted to number conditions 1–9.

For each subject, we constructed a trial-by-feature population matrix by concatenating the selected neural features across neurons. Neurons were included if they passed a number-selectivity threshold of *p* ≤ 0.125 in the single-neuron decoding analysis. For LLM, all valid embedding dimensions were used without any screening based on the selectivity of each dimension.

The population features were obtained from pooling at most 3 temporal LDA-component projections per neuron over all selective neurons for each subject. The feature matrix was z-scored across trials before computing geometry.

For each number condition *c*, we computed the empirical population center:

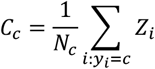

where *Z*_*i*_ is the population response on trial *i, y*_*i*_ is its number label, and *N*_*c*_ is the number of trials for number *c*. Sequential transition vectors were then defined as:

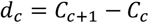

For each adjacent pair of transitions, we computed the transition angle:

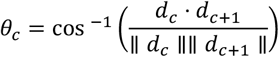

Angles were expressed in degrees and bounded between 0^°^and 180^°^. A one-dimensional ordered number line predicts transition angles near 0^°^, because all sequential transitions point in the same direction. A regular simplex predicts transition angles near 120^°^, because consecutive simplex edge directions are separated by 120^°^.

### Split-half empirical-noise procedure

Because trial noise can inflate transition angles even when the underlying class centers lie on a line, we estimated transition-angle distributions using split-half residual resampling. For each number condition, we decomposed each trial response into a condition center and a residual:

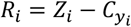

On each resampling iteration, residuals were sampled separately for each number condition and divided into two non-overlapping halves. The number of residuals per half was:

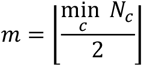

For each condition *c*, the sampled residuals were averaged within each half to obtain two residual-mean estimates, 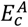 and 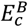. These residual means were then added back to the model centers, producing split-half noisy center estimates. For the empirical data:

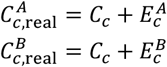

This procedure preserves the empirical class-specific trial noise while allowing the underlying center geometry to be replaced by matched model geometries.

Transition angles were computed across split halves rather than within the same split. Specifically, for each resampling iteration, we computed the angle between transition *c* from split A and transition *c* + 1from split B, and also the reverse comparison between transition *c*from split B and transition *c* + 1from split A:

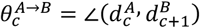

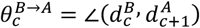

For nine number conditions, this yielded 2 × 7 = 14 transition-angle estimates per resampling iteration. The mean transition angle for an iteration was the average across these 14 angles. We repeated this procedure 1000 times per subject.

### Noisy log-number-line control

To construct the number-line control, empirical number centers were shifted onto a fitted one-dimensional number line while preserving the empirical residual noise. For the log-number-line control, each number condition *c* was assigned the scalar coordinate:

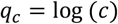

The coordinate vector was standardized across number conditions, and each neural feature was regressed onto this coordinate:

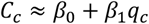

This produced fitted number-line centers:

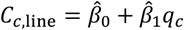

These fitted centers lie on a single ordered axis in population space. The same split-half residual means used for the empirical data were added to the number-line centers:

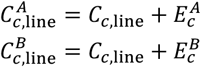

Thus, the noisy number-line control preserved the empirical trial-to-trial variability and class-specific sampling noise, but forced the underlying condition centers to lie along a fitted ordered line. Transition angles were then computed using the same cross-split procedure as for the empirical data.

### Matched simplex control

To construct the matched simplex control, we generated a full-rank simplex-like template whose class-specific radii matched the empirical distance of each number center from the global centroid. For each empirical center, the radial distance was defined as:

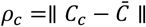

To avoid collapse when a radial distance was near zero, each radius was lower bounded by 5% of the root-mean-square radius.

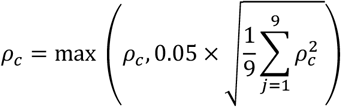

These radii were used to construct a diagonal, eye-like template with one class initially assigned to each axis. The template was then centered across classes, reduced to *S* − 1 dimensions using singular value decomposition, scaled to match the total empirical centroid variance, and embedded into the empirical neural feature space using the orthonormal orientation that best aligned the template with the empirical condition centers. This produced a full-rank simplex-like reference with empirical radial scale while preserving high-dimensional separation among number centers. Transition angles were then computed using the same cross-split procedure.

### Numerical-distance analysis of representational geometry

We tested whether representational distance increased with numerical distance, as predicted by an ordered number-line representation. This analysis was performed separately for four formats: nonsymbolic dot numerosities during the dot-counting stimulus epoch, Arabic numerals during the arithmetic task, Arabic-numeral feedback during the dot-counting task, and GPT-J embeddings of numerical stimuli.

For the three neural formats, trial-level population representations were constructed from the temporal LDA features described above. For each subject and task epoch, neurons were included if they passed the lenient number-selectivity threshold used for geometry analyses: *p* ≤ 0.125.

For each included neuron, we projected trials onto the neuron’s optimized temporal LDA components. These components were obtained from the best temporal window and shrinkage parameter identified in the single-neuron decoding analysis. Up to the first three LDA components were retained for each neuron. If a neuron’s optimized representation contained only one feature, that feature was passed through directly. Components from all included neurons were concatenated to form a trial-by-feature population matrix. Features were z-scored across trials before computing distances.

For the dot-stimulus format, trials were labeled by the number of dots shown. For the dot-feedback format, trials were labeled by the Arabic numeral shown during the feedback epoch. For the arithmetic-numeral format, operand 1 and operand 2 presentations were pooled as described above; each operand presentation was labeled only by its numerical value.For GPT-J, we used the extracted embedding vectors described in later sections after z-scoring.

For each subject and format, we computed the centroid of each number condition in the corresponding feature space. Let

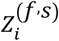

denote the feature vector for trial *i*from format *f*and subject *s*, and let *y*_*i*_be its number label. The centroid for number *c* was:

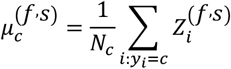

Representational distance between two numbers *c* and *d* was defined as the Euclidean distance between their centroids:

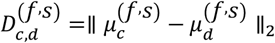

The corresponding numerical distance was:

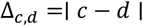

We used centroid distances rather than average pairwise trial-to-trial distances, because the analysis asked whether the stable number-condition centers were arranged according to numerical magnitude. For each subject and format, the upper triangle of the representational distance matrix was extracted. The diagonal entries, corresponding to zero numerical distance, were excluded from the mixed-effects model unless otherwise stated. Therefore, the primary model tested distances between distinct number identities, with numerical distances ranging from 1 to 8.

To test for a numerical-distance effect, we fit a linear mixed-effects model separately for each format:

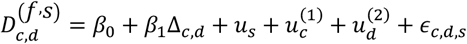

Here, 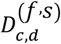 is the representational distance between numbers *c* and *d*, Δ_*c,d*_ is their absolute numerical distance, *u*_*s*_ is a random intercept for subject, and 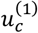 and 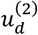 are random intercepts for the two number identities in the matrix entry. In MATLAB notation, the model was:

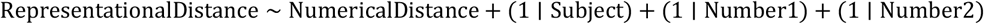

The random effects for Number1 and Number2 accounted for the fact that entries in a representational distance matrix are not independent, because each number appears in multiple pairwise distances. The fixed-effect slope *β*_1_ tested whether representational distance increased as numerical distance increased. A positive significant slope would be more consistent with a number-line-like metric structure.

For visualization, distances were also averaged by numerical distance within each subject. Subject-level distance curves were plotted in gray, and group means were plotted with s.e.m. across observations at each numerical distance. The fixed-effect prediction from the mixed-effects model was overlaid for visualization only. The statistical inference was based on the mixed-effects model fit to the unaveraged distance-matrix entries.

### Ordered number-line variance decomposition

To test whether the number manifold contained a privileged ordered number-line axis, we performed a variance-decomposition analysis on the same number-evoked population responses used for the Figure 2 geometry analyses. Analyses were performed separately for each subject and were restricted to numbers 1–9. Neurons were included if they passed the lenient number-selectivity criterion used for geometry analyses: *p* ≤ 0.125.

For each included neuron, we projected trials onto the neuron’s optimized temporal LDA components. These components were obtained from the best temporal window and shrinkage parameter identified in the single-neuron decoding analysis. Up to the first three LDA components were retained for each neuron. If a neuron’s optimized representation contained only one feature, that feature was passed through directly. Components from all included neurons were concatenated to form a trial-by-feature population matrix. When analyzing LLM embeddings, first 50 principal components were taken for subsequent analysis.

All variance percentages below are expressed relative to the total single-trial variance in this analysis space.

Let

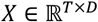

denote the resulting population response matrix for one subject, where *T* is the number of trials and *D* is the number of retained features. Each trial *i* had a number label

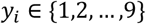

Before the variance decomposition, the trial-wise population matrix was mean-centered across trials:

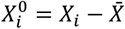

For each number condition *c*, we computed the condition mean response:

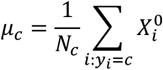

We then replaced each single-trial response with the mean response for its corresponding number condition:

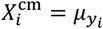

This condition-mean representation removes trial-by-trial variability while preserving stable number-dependent differences in the population response. The amount of reliable number-condition structure was quantified as the percentage of total trial-level variance explained by this condition-mean matrix:

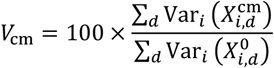

We next asked how much of this reliable condition-mean structure lay along a continuous ordered number-line axis. We tested both a linear number line and a logarithmic number line. For the linear model, the target coordinate for number *c* was

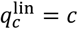

For the logarithmic model, the target coordinate was

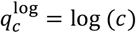

For each target coordinate system, we fit a ridge-regression axis from single-trial population responses to the corresponding number coordinate. The ridge penalty was set to

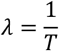

where *T* is the number of trials. The fitted weight vector was normalized to unit length before computing variance along the axis.

For a given number-line coordinate *q*, the fitted number-line axis can be written as

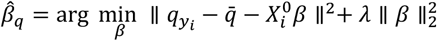

followed by normalization:

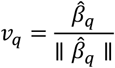

We then projected the condition-mean responses onto this fitted axis and quantified the fraction of total trial-level variance captured by the number-line component:

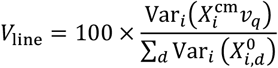

The residual reliable condition-mean variance not captured by the number-line component was then defined as

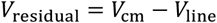

This calculation was performed separately for the linear and logarithmic number-line axes. For each subject, we therefore obtained a linear number-line variance component, a linear residual condition-mean component, a log-number-line variance component, and a log residual condition-mean component.

Across subjects, the number-line component was compared with the corresponding residual condition-mean component using a two-sided paired *t*-test. Linear and logarithmic number-line decompositions were tested separately. Values are reported as mean ± SEM across subjects.

### All-order single-trial number-line rank analysis

As a complementary test, we asked whether the canonical 1–9 ordering was privileged relative to all other possible one-dimensional orderings of the same nine number identities. This analysis used the same subject-level population matrix *X*and number labels *y*_*i*_as the variance-decomposition analysis.

For each subject, we evaluated all

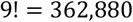

possible assignments of the nine number identities to positions on a one-dimensional ordered line. Each permutation *π* defined a candidate ordered line. For the linear analysis, number identity *c*was assigned the permuted coordinate

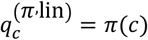

For the logarithmic analysis, number identity *c* was assigned

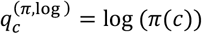

The canonical number line corresponded to the identity ordering:

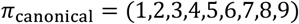

For each candidate ordering, we trained a ridge-regression model to predict the corresponding one-dimensional coordinate from single-trial population activity. Performance was estimated using stratified 3-fold cross-validation. If any number condition had fewer than three trials, the number of folds was reduced to the minimum class count, with a minimum requirement of at least two trials per number condition.

Within each cross-validation fold, features were z-scored using the training trials only. The training-set mean and standard deviation were then applied to the held-out test trials. The ridge penalty was again set to

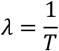

For each held-out fold, the model predicted the one-dimensional coordinate of each test trial. Predictions were concatenated across folds. Decoding performance for a given ordering was quantified as the Pearson correlation between the predicted and target coordinates across all held-out trials:

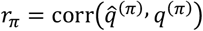

For each subject, the percentile rank of the canonical number line was computed relative to the full distribution of correlations obtained from all 9 ! possible orderings:

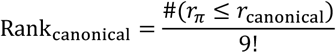

A rank of 0.5 indicates that the canonical ordering performed no better than a typical arbitrary ordering. A rank of 1 indicates that the canonical ordering was the best possible ordered line among all 9 ! permutations.

Canonical ranks were computed separately for the linear and logarithmic coordinate systems. Across subjects, we tested whether the canonical rank was above chance using a one-sided Wilcoxon signed-rank test against 0.5. We also tested whether the canonical rank remained below the best possible ordering using a one-sided Wilcoxon signed-rank test against 1.0. Linear and logarithmic canonical ranks were compared with a paired Wilcoxon signed-rank test.

### Shattering dimensionality analysis

We quantified the linear separability of number representations using a shattering-dimensionality analysis (Bernardi et al., 2020; Posani et al., 2025). Analyses were performed separately for each subject. Unless otherwise stated, analyses were restricted to number conditions 1–9.

For each subject, we constructed a trial-by-feature population matrix from MTL neurons. Neurons were included if they passed a lenient number-selectivity threshold of *p* ≤ 0.125 in the single-neuron number-decoding analysis. For each included neuron, we used the temporal LDA projection described above. Specifically, the previously optimized single-neuron LDA model was used to project each trial onto that neuron’s number-discriminating temporal components. Up to the first three temporal components were retained per neuron. If a neuron had only one feature, that feature was passed through directly. The retained components from all included neurons were concatenated to form the full population representation.

No PCA was applied to the neural data before computing the full shattering-dimensionality score. PCA was used only for the explicit low-dimensional control analyses described below. For LLM embedding analyses, the embedding matrix was first reduced to PC1–50 before applying the same shattering pipeline so that the number of features does not overwhelm the number of trials; this PC1–50 preprocessing was not applied to neural population data.

Number classes were included if they had at least two trials for that subject. Let *P* denote the number of valid classes for a subject. In the standard 1–9 analysis, *P* = 9. Trials were labeled by their number condition, and the valid number labels were mapped to class indices 1,2, …, *P*.

### All-dichotomy shattering score (simplex shattering dimensionality)

For each subject, we evaluated binary decoding across all unique dichotomies(Rigotti et al., 2013) of the valid number classes. Rather than restricting the analysis to balanced half-splits (Bernardi et al., 2020; Posani et al., 2025). We considered splits of all sizes from one class versus the remaining classes up to

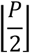

classes versus the remaining classes. Thus, for *P* = 9, the total number of dichotomies was

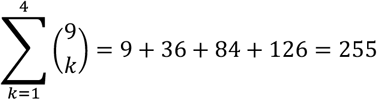

For a given dichotomy, one subset of number classes was assigned to binary class 0, and the complementary subset was assigned to binary class 1. The sign of this assignment was arbitrary because performance was measured as classification accuracy.

For each dichotomy, all trials belonging to numbers on one side of the split were pooled together, and all trials belonging to numbers on the other side were pooled together. We did not equalize trial counts separately for each number. Instead, within each dichotomy and each resampling iteration, we balanced only the total number of trials assigned to the two binary classes by randomly downsampling (160 times) the larger binary side to match the smaller side. This produced a balanced binary classification problem for each dichotomy, with chance performance equal to 0.5.

### Cross-validated dichotomy decoding

Each balanced binary classification problem was decoded using a ridge-regularized linear support-vector classifier. Classification was performed with MATLAB’s fitclinear using a linear SVM loss and ridge regularization.

For each dichotomy and resampling iteration, the balanced trial set was divided into stratified cross-validation folds. The requested maximum number of folds was 5, but the actual fold number was reduced when trial counts were limited:

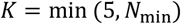

where *N*_min_ is the number of trials in the smaller binary class after balancing. Within each fold, features were z-scored using only the training trials, and the same training-set mean and standard deviation were applied to the held-out test trials. The decoding accuracy for one dichotomy was defined as the mean held-out accuracy across folds. Because the two binary classes were balanced before cross-validation, ordinary accuracy was equivalent to balanced accuracy for this analysis.

To stabilize the estimate against random trial downsampling, the full dichotomy-decoding procedure was repeated 160 times. For each dichotomy, the decoding accuracy was averaged across these 160 repetitions. The simplex shattering-dimensionality score, denoted SSD, was then defined as the mean decoding accuracy across all dichotomies:

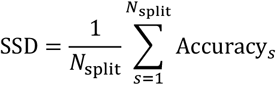

where *N*_split_ = 255 for the full nine-number analysis.

### Label-shuffle control

To assess whether shattering performance exceeded chance, we generated a label-shuffle null distribution for each subject. On each shuffle iteration, number labels were randomly permuted across trials while preserving the neural data and the total label counts. The same dichotomy construction, trial balancing, cross-validation, and classifier-fitting procedure was then repeated. The shuffle analysis used the same number of iterations as the observed analysis: *N*_shuffle_ = 160.

For each shuffle iteration, we computed the mean accuracy across all dichotomies, yielding a subject-specific null distribution of SSD values. A one-sided empirical p-value was computed as

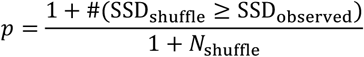

A subject’s shattering score was considered above chance if its observed SSD exceeded the shuffled-label distribution at *p* < 0.05.

### Regressing out magnitude-related structure

To test whether high shattering performance could be explained by a simple number-magnitude axis, we repeated the analysis after removing magnitude-related variance from the population representation. For each feature dimension, activity was regressed against number magnitude across trials. In the final implementation, the magnitude regressor was log-transformed number:

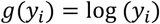

For each feature *d*, we fit

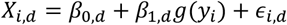

The residuals

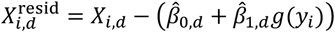

were used as the magnitude-regressed population representation. We then recomputed SSD using the same all-dichotomy decoding procedure. Group-level significance of the magnitude-regressed SSD was assessed across subjects by testing whether the residual SSD remained above chance level (one dimensional), SSD > 0.5.

### Low-dimensional PCA controls

To estimate the minimum dimensionality needed to account for the observed shattering performance, we projected the neural population into low-dimensional subspaces and recomputed SSD. PCA axes were computed from number-condition centers, not from the full single-trial covariance matrix.

For each subject, the mean population response for each number condition was computed as

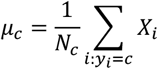

The matrix of condition means was centered across number classes and decomposed with PCA. We then projected each single-trial population response onto the first *d* condition-mean PCs:

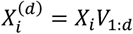

where *V*_1:*d*_contains the first *d* PCA axes. We recomputed SSD for

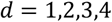

These PCA-projected analyses were used only as dimensionality controls. The full neural SSD reported in the main analysis was computed from the original concatenated neural feature space and was not PCA-reduced.

### Cross-condition generalization performance

We used cross-condition generalization performance (CCGP) to test whether number-related population activity contained latent categorical axes that generalized across held-out number conditions, following the logic of abstract-variable decoding analyses (Bernardi et al., 2020).

For each subject, trials were labeled by the number associated with that trial and analyses were restricted to numbers 1–9. Number classes with fewer than two trials were excluded for that subject. If a predefined dichotomy contained a missing or under-sampled number for a given subject, the dichotomy was intersected with the subject’s valid classes. A subject–dichotomy combination was analyzed only if at least two classes remained on each side of the split.

Population features were constructed from the single-neuron temporal LDA projections described above. For each neuron, the previously optimized temporal LDA model was used to project each trial onto that neuron’s number-discriminating temporal components. Up to the first three temporal components were retained per neuron. If a neuron’s optimized representation contained only one feature, that feature was passed through directly. Neurons were included if they passed a lenient number-selectivity threshold of *p* ≤ 0.125 in the single-neuron number-decoding analysis. All retained components from included neurons were concatenated to form a trial-by-feature population matrix. LLM embedding dimensions were not shortlisted by selectivity. First 50 principal components were used for LLM embedding.

### Number dichotomies

We tested 11 predefined dichotomies of the numbers 1–9. The sign of the binary label was arbitrary; for each split, one set was assigned to class −1 and the complementary set to class +1. The implemented dichotomies were:

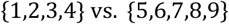

for small versus large numbers;

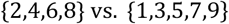

for even versus odd numbers;

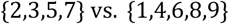

for prime versus non-prime numbers;

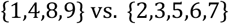

for perfect squares or cubes;

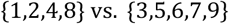

for powers of two;

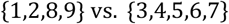

for numbers far from the center of the range;

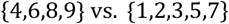

for a visual closed-shape grouping;

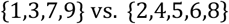

for keypad-corner numbers;

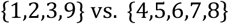

for Roman-numeral forms without versus with the letter V;

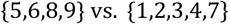

for English number words containing the letter i; and

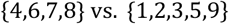

for English number words ending in a consonant versus a vowel.

### Leave-one-number-pair-out CCGP

For each dichotomy, we used a leave-one-number-from-each-side-out procedure. On each split, one number from the −1 side and one number from the +1 side were held out as test conditions. The classifier was trained on all remaining numbers in the dichotomy and tested on the held-out pair. For example, for the small-versus-large split, one fold could train on:

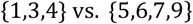

and test whether this axis generalized to:

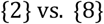

Thus, for a split with *n*_−_ classes on one side and *n*_+_ classes on the other side, the number of held-out train/test splits was:

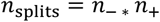

For the standard 4-versus-5 dichotomies used here, this yielded:

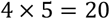

held-out splits.

Within each held-out split, all trials from the training numbers were pooled according to their binary category label. To prevent unequal trial counts from biasing the classifier, the larger binary training class was randomly downsampled to match the smaller training class. This balancing was repeated 16 times per held-out split, and decoding accuracy was averaged across these repetitions.

For each balancing repetition, features were z-scored using the training data only, and the same centering and scaling parameters were then applied to the held-out test data. A linear classifier with logistic loss and ridge regularization was fit to the training trials using MATLAB’s fitclinear implementation. The regularization parameter was set using the classifier’s automatic setting.

Test performance was measured using balanced accuracy:

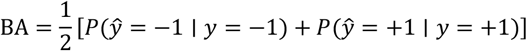

Balanced accuracy was used so that unequal numbers of held-out trials from the two test numbers did not bias the CCGP estimate. Chance performance was therefore:

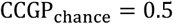

The CCGP value for a given subject and dichotomy was defined as the mean balanced accuracy across all held-out number-pair splits and all training-set balancing repetitions:

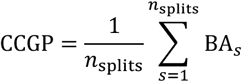

### Null distributions and subject-level significance

We computed a neuron-shuffle geometric null (Bernardi et al., 2020) for each subject and dichotomy. For each number class, the population-feature columns were randomly permuted independently before constructing the training and test sets. This preserved the marginal activity distribution within each number class but disrupted the consistent identity of neural features across number conditions. This null tested whether apparent cross-condition generalization could arise from class-wise activity distributions alone, without a stable population axis shared across numbers. Each null distribution used 100 shuffles. For each null, a one-sided empirical p-value was computed as:

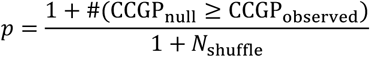

For the subject-level significance marks shown in Figure 2I, a subject was counted as showing significant generalization for a dichotomy if the observed CCGP exceeded the neuron-shuffle null at *p* < 0.05. The number shown on each bar indicates the number of subjects with significant neuron-shuffle-corrected CCGP for that dichotomy.

### GPT-J embedding extraction for arithmetic stimuli

We extracted language-model embeddings from GPT-J-6B using the Hugging Face Transformers implementation of EleutherAI/gpt-j-6b (Wang and Komatsuzaki, 2021). The model was loaded with output_hidden_states = True, half-precision weights. The model was set to evaluation mode, and all forward passes were performed with gradients disabled using torch.no_grad(). No fine-tuning or text generation was performed; embeddings were extracted only from fixed arithmetic prompt strings.

For each arithmetic trial, we used the first operand, second operand, operation sign, and operation-position information. Downstream analyses used only three semantic slots:

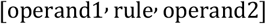

The equality token and answer token were not analyzed. Although the implementation was capable of constructing full equation strings, the embeddings used in the analyses were extracted from the first three semantic elements only. Because GPT-J is an autoregressive model, the hidden state for a token depends only on preceding tokens and the current token. Therefore, embeddings for operand 1, the rule, and operand 2 are unchanged by later equality or answer tokens. For this reason, embeddings corresponding to the participant’s given answer or the task-defined correct answer were not included in the analyses.

Numeric stimuli were represented in two formats. In the symbolic format, numbers were written as Arabic numerals, including signed values when present. In the language format, numbers were converted to English words using an integer-to-word conversion routine. Negative numbers were written with the prefix “negative,” for example −5 was written as “negative five.”

We generated four prompt families for each trial.

First, we generated a symbolic canonical prompt in standard operand-rule-operand order:

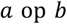

For example, a trial with operands 3 and 4 and addition was represented as:

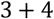

Second, we generated a symbolic operation-position-conditioned prompt. When the operation sign appeared before the operands in the task, the prompt used a prefix-like form:

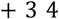

When the operation sign did not appear first, the prompt used a postfix-like form:

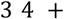

The model saw the tokens in this surface order, but extracted embeddings were stored in fixed semantic order as operand 1, rule, and operand 2.

Third, we generated a language canonical prompt. Addition trials used the operator words “plus” and “add,” and subtraction trials used “minus” and “subtract.” For example, an addition trial was represented as both:

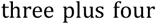

and

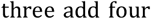

Embeddings from the two operator-word variants were averaged elementwise to obtain a single language canonical representation for that trial. The same procedure was used for subtraction trials.

Fourth, we generated a language operation-position-conditioned prompt. When the operation sign appeared first, the prompt used a prefix-like language form, for example:

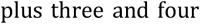

or

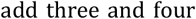

When the operation sign did not appear first, the prompt used a postfix-like language form, for example:

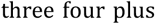

The support word “and” was included to make prefix-like language prompts more natural, but it was not included in the extracted operand-2 embedding whenever token offsets allowed the operand word to be isolated.

The four retained GPT-J embedding sets were therefore:

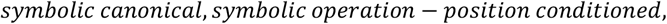

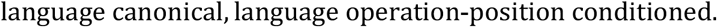

For each set, embeddings were extracted incrementally to respect the causal structure of GPT-J. For each prompt, the running text was initialized as an empty string. Prompt elements were then appended one at a time. After each append operation, the current prefix was tokenized and passed through GPT-J. The hidden states corresponding to the newly appended target element were then extracted. Thus, the operand 1 embedding was extracted from the prefix containing operand 1 alone, the rule embedding was extracted after appending the operator to the current prefix, and the operand 2 embedding was extracted after appending the second operand.

For each target element, tokenizer character offsets were used to identify the tokens corresponding to the exact appended target span. When a target element was split into multiple tokens, the hidden states of those tokens were averaged. This procedure was used for multi-token number words such as “negative five” as well as for any multi-token symbolic strings.

For GPT-J-6B, hidden states were extracted from the embedding layer and all transformer layers, yielding 29 hidden-state levels with 4096 dimensions per level. For each trial and prompt family, the retained embedding tensor used for analysis therefore had dimensions:

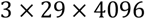

corresponding to semantic slot, hidden-state level, and embedding dimension. For downstream representational analyses, embeddings were taken from 26^th^ layer.

For each embedding set(format), embeddings were further normalized separately for the two operand positions. Let

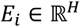

denote the GPT-J embedding vector for operand-level row *i*, and let

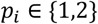

denote whether that row corresponded to operand 1 or operand 2. For each operand position *p*, we first subtracted the operand-position-specific mean embedding:

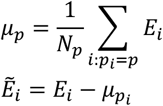

This step removed fixed embedding offsets associated with whether a number appeared as the first or second operand, so that the embedding emphasized trial-specific numerical representational variation rather than a global operand-position difference.

We then removed embedding dimensions with unusually large variance across the centered operand-level rows. For each embedding dimension *d*, variance was computed as

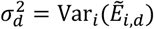

A dimension was excluded if

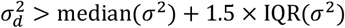

where the median and interquartile range were computed across embedding dimensions. This robust exclusion step was included because information in contextual Transformer embeddings can be dominated by a small number of high-variance or high-magnitude dimensions that do not necessarily reflect the stimulus variable of interest. Prior work has shown that such “rogue” or outlier dimensions can dominate cosine and Euclidean similarity, can mismatch dimensions that are important for model behavior, and can correlate with non-semantic factors such as token position, punctuation, or token frequency (Timkey and van Schijndel, 2021; Puccetti et al., 2022).

Finally, each operand-position group was scaled by its mean row-wise vector norm. For operand position *p*, we computed

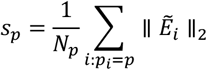

and the final normalized embedding was

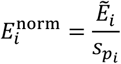

This scaling matched the average embedding magnitude across the two operand positions while preserving the relative geometry among trials within each operand position.

### Trial-level comparison between MTL arithmetic geometry and GPT-J geometry

We compared trial-level representational geometry during the arithmetic task with two candidate model geometries: a numerical-distance model and a GPT-J embedding-distance model. Analyses were performed separately for each subject.

The analysis used the cue-expanded arithmetic representation. In this representation, each arithmetic formula trial contributed two number-trial rows: one row for the first operand and one row for the second operand. Thus, if a subject completed *N* arithmetic trials, the representational matrix contained up to 2*N* operand-level rows.

We used the LDA-based neural representation described above. Briefly, for each neuron, the previously optimized single-neuron temporal LDA model was used to project each operand-level trial onto number-discriminating temporal components. LDA axes were fit only using valid rows and the number classes available for fitting, but after fitting, all operand-level rows were projected, including rows from low-count number classes that were not used to fit the LDA axes. Up to the first three components per neuron were retained. Only tuned neurons were used for the main analysis. A neuron was included if it passed the lenient tuning criterion used for the geometry analyses: *p* ≤ 0.125 and its decoding accuracy exceeded chance for the fitted class set. The retained LDA components were concatenated across neurons to form a trial-by-feature matrix:

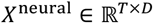

where *T* is the number of trials and *D* is the number of retained neural features. Before computing distances, each neural feature was mean-centered across trials:

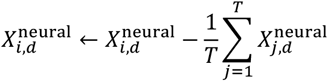

For each subject, the neural representational distance matrix was computed using Euclidean distance between individual operand-level trial responses:

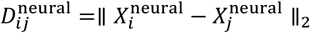

The diagonal was excluded, and only the upper triangle of the RDM was used for model comparison.

### Numerical-distance model RDM

For each operand-level trial *i*, let *n*_*i*_ denote the displayed operand value associated with that row. We constructed numerical model RDMs directly from these trial-wise number values. We used an asinh-transformed, log-like numerical-distance model:

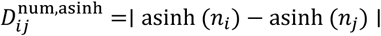

note:

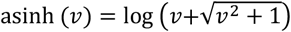

### GPT-J embedding RDM

For GPT-J embeddings, we used operand-level embeddings matched to the same trial rows as the neural arithmetic representation. Each arithmetic trial contributed one row for operand 1 and one row for operand 2. GPT-J embeddings were taken from layer 26 and preprocessed as described in the previous section and were compared with neural RDMs using Euclidean distance.

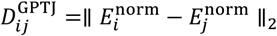

The vectorized upper triangle of this GPT-J RDM was correlated with the vectorized upper triangle of the neural RDM using Pearson correlation. For comparison, we also constructed numerical-distance RDMs from the operand values and computed their correlations with the neural RDM using the same procedure.

### Arithmetic operand-to-result transformation analysis

We tested whether arithmetic result-related population activity could be predicted from neural representations of the two operands and the arithmetic operation. Analyses were performed separately for each subject. Operand representations were extracted from the response-adjusted arithmetic configuration described above. Trials with missing participant-entered answers were excluded. Trials were also excluded if their reaction time RT exceeded:

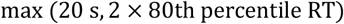

The behavioral target was the participant’s entered answer, not necessarily the objectively correct answer.

### Thinking-period result representation and answer-decoder grid search

Before testing operand-to-result transformations, we first fit subject-specific decoders to determine whether MTL activity during the thinking period predicted the answer the participant would enter. The thinking-period activity of each trial was evenly divided into 12 non-overlapping segments. For most subjects, bins 1–12 were used. For subject YFJ, we used bins 7–12 for the arithmetic thinking-period decoder due to a consistently longer thinking period against the rest subject population.

For each candidate result representation, the selected thinking-period bins were further divided into *n*_windows_ contiguous temporal windows. We searched:

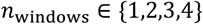

Within each window, spike counts were summed for each neuron. The summed counts were divided by the corresponding fraction of the trial’s reaction time,

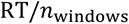

yielding a firing-rate vector for each temporal window. Vectors from all windows were concatenated, so that the dimensionality of the result representation was:

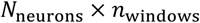

Thus, the result representation *Y*_res_ was not fixed a priori as a single firing-rate vector or as all 12 bins separately. Instead, the number of temporal windows used to serialize thinking-period activity was selected by grid search jointly with the answer-decoder model.

For each subject and each candidate *n*_windows_, we evaluated three families of answer decoders using 20-fold cross-validation. The target variable was the participant’s entered answer.

The first family was a linear model implemented with fitrlinear. We searched two learners,

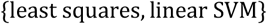

two regularization types,

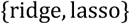

and regularization strengths:

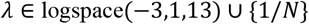

where *N* is the number of trials. The logspace function samples 1e-3 to 1e1 with 13 log-spaced samples.

The second family was an RBF-kernel support-vector regression model implemented with fitrsvm. We searched:

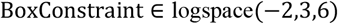

and

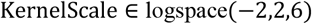

together with MATLAB’s default RBF-SVM configuration.

The third family was an ensemble regression model implemented with fitrensemble using tree learners.

Ensemble hyperparameters were optimized by MATLAB’s automatic hyperparameter optimization under the same cross-validation partition. The optimized parameters included the ensemble method, number of learning cycles, minimum tree leaf size, and learning rate when applicable.

For the linear and SVM decoders, features were z-scored within each training fold, and the training-fold mean and standard deviation were applied to the held-out fold. Ensemble decoders were fit using the raw result-representation features. For every model and window configuration, decoding performance was quantified using cross-validated *R*^2^:

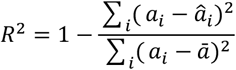

where *a*_*i*_ is the participant’s entered answer, â_*i*_ is the predicted answer, and ā is the mean entered answer across evaluated trials. Mean absolute error was also computed.

For each subject, the best decoder was chosen as the model and *n*_windows_ configuration with the highest cross-validated *R*^2^. The resulting best representation matrix was designated as *Y*_res_ and used as the target result representation in the operand-to-result transformation analysis below.

To assess whether thinking-period answer decoding exceeded chance, we repeated the best-model procedure using shuffled answer labels. The same selected decoder family, hyperparameters, window count, and cross-validation partition were used. We used 200 label shuffles. A one-sided permutation p-value was computed as:

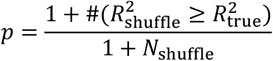

For subject YFU, predicted answers were lower-bounded at zero before computing *R*^2^, rounded accuracy, and mean absolute error since the subject never inputted negative number answers.

### Operand representations

Operand representations were defined from the single-neuron temporal LDA projections described above. We further restricted the operand representation to neurons that significantly encoded operand value in the response-adjusted operand analysis: *p* < 0.05. For each retained neuron, operand 1 and operand 2 trials were projected into a shared LDA-derived feature space. This was done by concatenating operand-1 and operand-2 trials before projection and then splitting the projected features back into matched operand-1 and operand-2 matrices. This ensured that both operands were represented in the same component coordinate system.

Let *X*_*op*1,*i*_ and *X*_*op*2,*i*_ denote the operand-1 and operand-2 feature vectors for trial *i*. Each vector contained the concatenated temporal-component coordinates across all retained operand-tuned neurons.

### Operand-to-result model variants

We compared five operand-to-result transformation models. All models predicted the thinking-period result representation *Y*_res_ from operand features using multi-input, multi-output ridge regression.

The Op1-only model, used only the first operand:

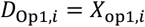

The Op2-only model, used only the second operand:

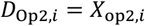

The Op12 model, used both operands but ignored the operation rule:

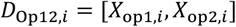

The flexible arithmetic model used operand 1, operand 2, and a rule-gated operand-2 term. The rule variable was defined as:

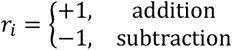

The flexible design matrix was:

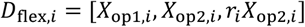

Thus, in the implemented flexible model, the contribution of operand 2 was allowed to depend on the arithmetic operation.

The commutative model used shared operand weights during addition while allowing a distinct operand-2 contribution during subtraction. We defined:

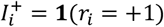

and

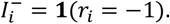

The commutative design matrix was:

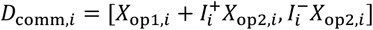

On addition trials, this reduces to:

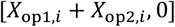

so operand 1 and operand 2 share the same weight matrix. On subtraction trials, it reduces to:

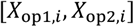

allowing operand 1 and operand 2 to contribute differently.

### Nested fitting of operand-to-result transformations

For each transformation model, the design matrix *D* was mapped to *Y*_res_ using ridge regression. Performance was estimated using 10-fold outer cross-validation. Within each outer training fold, the ridge penalty was selected using 5-fold inner cross-validation.

The ridge penalty grid was:

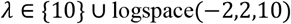

Within each training fold, the design-matrix features were z-scored using training-set statistics only, and the same mean and standard deviation were applied to held-out trials. The result-representation matrix was mean-centered in the training set. Ridge weights were estimated as:

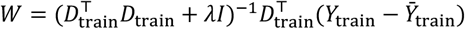

Predicted result representations for held-out trials were computed as:

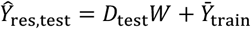

Inner-fold hyperparameter selection was decoder-driven. For each candidate *λ*, the predicted validation-fold result representation was passed through the fold-refit answer decoder described below. The selected *λ* was the value that maximized validation evidence. For numeric answer decoding, this evidence measure was negative absolute answer error. The final reported metric, however, was cross-validated *R*^2^ between predicted and actual participant-entered answers.

### Fold-wise refitting of the answer decoder

The estimated result representation Ŷ_res_ was not passed through a single fixed answer decoder trained on all trials. Instead, the answer decoder was refit within each outer transformation fold.

For each subject, the best result-decoder family, temporal window count, and hyperparameters were obtained from the grid-search result. Within each outer fold of the operand-to-result analysis, a decoder with the same selected model type and hyperparameters was fit using the true thinking-period result representation *Y*_res_ and participant-entered answers from the training trials only. This fold-refit decoder was then applied to the synthetic held-out result representation Ŷ_res_ to predict the participant’s entered answer on held-out trials.

The same fold-refit decoder was also applied to the true held-out thinking-period representation *Y*_res_, providing a matched benchmark for how well the subject’s actual MTL thinking-period activity predicted upcoming answers.

For linear and SVM result decoders, the refit decoder used training-fold z-scoring. For ensemble decoders, the selected ensemble method, number of learning cycles, minimum leaf size, and learning rate were reused. When the selected ensemble minimum leaf size exceeded 100, the fold-refit tree template capped this parameter at 100.

Predicted answers were concatenated across outer folds. Model performance was quantified as:

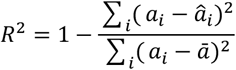

For YFU, the same nonnegative prediction cap used in the thinking-period result decoder was also applied

### Model comparison

For each subject, the best full arithmetic model was defined as the better of the flexible and commutative models:

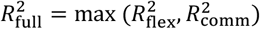

The best baseline model was defined as the best of the three ablated models:

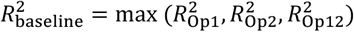

### Number-line operand compression controls

To test whether arithmetic transformations were better explained by first compressing operand representations into a one-dimensional number-line variable, we repeated the operand-to-result analysis after supervised one-dimensional operand projection.

This projection was fit separately within each cross-validation fold. Within a training fold, operand 1 and operand 2 feature matrices were concatenated:

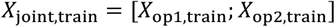

and the corresponding operand values were concatenated:

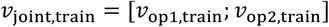

For the linear number-line control, a linear model was fit from the high-dimensional operand representation to signed operand value:

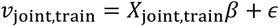

The fitted model was then applied to both training and held-out operand features, replacing *X*_*op*1_and *X*_*op*2_with one-dimensional predicted operand values.

For the log-like number-line control, the same fold-local procedure was used, but the regression target was:

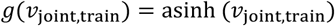

The inverse hyperbolic sine was used because it provides a logarithmic-like compression that remains defined for zero and negative operand values. Specifically,

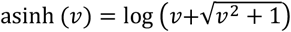

For large positive values,

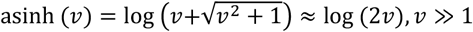

For large negative values, because asinh (*v*) is an odd function,

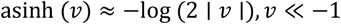

After this one-dimensional operand compression, the same five operand-to-result models were fit and evaluated. For each subject, the best full model from the original high-dimensional operand representation was compared with the best full model after linear operand compression and after asinh/log-like operand compression. These comparisons used paired Wilcoxon signed-rank tests across subjects.

### Thinking-period result decodability and behavioral correlations

To quantify whether MTL thinking-period activity contained an upcoming answer representation, we used each subject’s best result decoder from the thinking-period grid-search analysis. The decoder target was the participant’s entered answer. For each subject, we retained the selected model family, selected hyperparameters, cross-validated R^2^, and label-shuffle p value

Behavioral accuracy was computed as:

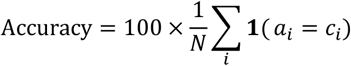

where *a*_*i*_ is the participant’s entered answer and *c*_*i*_ is the task-defined correct answer. Mean absolute error was computed as:

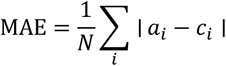

Trials with missing entered answers were excluded. Across subjects, result decodability was related to behavior by correlating each subject’s thinking-period result-decoding *R*^2^ with behavioral accuracy and MAE. Pearson correlation was used for statistical inference. A linear fit was used for visualization of the accuracy relationship. For the MAE relationship, an exponential curve was used for visualization, but the reported p-value was based on the Pearson correlation between behavioral MAE and result-decoding *R*^2^.

